# Insights into adipokinetic hormone/corazonin-related peptide receptor specificity and key residues for its activation in the human disease vector *Aedes aegypti* mosquito

**DOI:** 10.1101/2023.05.16.541050

**Authors:** Salwa Afifi, Jean-Paul Paluzzi

## Abstract

Adipokinetic hormone/corazonin-related peptide (ACP) and adipokinetic hormone (AKH) are two neuropeptides that demonstrate homology to the vertebrate gonadotropin-releasing hormone (GnRH). Despite the structural similarity and the close evolutionary relationship between the ACP and the AKH, their signaling systems function independently. To date, the role of ACP and its receptor (ACPR) remains unclear in the *Aedes aegypti* mosquito. Structure-activity relationships (SARs) are often carried out on peptide ligands to determine critical residues for bioactivity and receptor activation; however, residues and features necessary for ligand binding and specificity in the receptors themselves are less studied. Herein, this study focuses on the ACP and AKH signaling systems and examines structural features of their receptors critical for conferring activation and ligand specificity. Firstly, to determine the specific ACPR regions most critical for ligand fidelity and specificity, ACPR chimeras were created by singly replacing each of the three ACPR extracellular loops (ECLs) in their entirety and incorporating the corresponding ECLs from the AKH receptor (AKHR). Heterologous functional assays determined that the three ACPR ECL chimera receptors with complete replacement of full individual ECLs showed no response to either ACP or AKH. These results suggest that the complete replacement of each individual extracellular loop is detrimental to ligand binding and recognition. Secondly, through a more targeted approach, we aimed to determine specific residues critical for functional ligand-binding by substituting only select highly conserved residues within the three ECLs of the ACPR with those from the AKHR. Modifications of specific and highly conserved residues in these ACPR ECLs chimeras suggest that the third extracellular loop contains the most critical residues necessary for ACP binding and receptor activation. In addition, the combination of two selectively-modified ACPR ECLs demonstrated a significant decrease in response to ACP compared to the native ACPR response. Interestingly, combining all of the ACPR ECLs chimeras together resulted in a significant decrease in response to ACP compared to native ACPR. Relatedly, a significantly increased response to AKH was observed in the receptor chimera combining selected modifications in all three ECLs compared to native ACPR. Hence, the particular residues essential for ACP ligand interaction were identified due to the detrimental effect that occurred in ACPR activation after the selective modification of crucial residues localized within the three extracellular domains of the receptor. These data provide key insight into how these two closely related neuropeptidergic systems maintain functional independence in the mosquito *A. aegypti* as well as other insects.

## Introduction

Mosquitoes are infamously known as vectors of pathogens causing several diseases in humans and other vertebrates. The mosquito *Aedes aegypti* is a chief vector of viruses including dengue and yellow fever, chikungunya and Zika that are the causative agents of illnesses with great impact on human health globally (Kotsakiozi et al., 2017). Due to climate change and globalization, the geographic range of these diseases has increased; therefore, a deeper understanding of the regulation of physiological processes, including those governed by neuropeptide signaling systems, is essential for insight to develop novel compounds geared towards vector control.

Neuropeptides and their receptors play a central role in the regulation of a wide variety of physiological and behavioral processes during the life cycle of arthropods, such as feeding, growth, metabolism, reproduction, locomotion and development. They are also fundamental in the chemical cell-to-cell communication systems between different types of cells (Nässel, 2002; Nässel and Homberg, 2006; Barón et al., 2010; Nässel and Winther, 2010; Nässel and Zandawala, 2019). Most neuropeptide receptors are members of the G protein-coupled receptors (GPCRs) superfamily, whose seven transmembrane domains are connected by three extracellular loops (ECLs) and three intracellular loops, with the former participating in the binding and interaction with a specific ligand (Caers et al., 2012b; Pierce et al., 2002; Vanden Broeck, 1996; Wise, 2012; Zandawala et al., 2015a).

Adipokinetic hormone (AKH), corazonin (CRZ) and adipokinetic hormone/corazonin related peptide (ACP) are three neuropeptides that were found in invertebrates and considered homologous to the mammalian gonadotropin-releasing hormone (GnRH) (Hansen et al., 2010; Gäde et al., 2011; Roch et al., 2011; Li et al., 2016). The ACP receptor (ACPR) is activated by a neuropeptide very closely related to both AKH and CRZ and was hence known as ACP (Hansen et al., 2010). Despite the ACP signaling system being structurally intermediate between AKH and CRZ signaling systems, the functional role of ACP and its receptor in most arthropods remains unclear. Moreover, these three neuropeptides and their receptors are not always found together in all arthropods. For instance, while mosquitoes have all three neuropeptide signaling systems (AKH, ACP, and CRZ), CRZ is absent in the beetle *Tribolium castaneum*, and ACP is missing in the fruit fly *Drosophila melanogaster* (Hansen et al., 2010). Interestingly, the lack of ACP in *Drosophila* has restricted the potential use of molecular genetic tools that could examine its physiological functions (Hansen et al., 2010; Zandawala et al., 2015a).

Phylogenetic analyses have previously indicated the AKH system duplicated before the emergence of the phylum Arthropoda, leading to the ACP and AKH neuropeptide systems (Hansen et al., 2010). Owing to gene duplication in arthropod lineage, ACP and AKH signaling systems were considered to be paralogous (Hauser and Grimmelikhuijzen, 2014). However, various studies showed the high selectivity of the ACP, AKH, CRZ receptors responding only to their cognate ligands. Therefore, these three neuropeptides and their receptors are indeed considered to work independently (Hamoudi et al., 2016; Hansen et al., 2010; Oryan et al., 2018; Wahedi and Paluzzi, 2018; Zandawala et al., 2015a). *In vitro* studies demonstrated that high concentrations of AKH can activate the ACP receptor and *vice versa* in the silkworm *Bombyx mori* (Zhu et al., 2009; Shi et al., 2011). However, it is important to note that CRZ failed to activate AKH and ACP receptors (and *vice versa*) in *Anopheles gambiae* and *Rhodnius prolixus*, which suggested that the AKH and ACP are more related to each other compared to the CRZ system (Hansen et al., 2010; Zandawala et al., 2015a). Despite acting through independent receptors, some actions of ACP are shared with AKH in *A. aegypti* including decreasing glycogen levels in the female fat body and hypertrehalosaemic activity in males; however, only AKH had hypertrehalosaemic action in females (Afifi et al., 2023). Thus, the ACP and AKH signaling systems are excellent examples of receptor-ligand co-evolution (Caers et al., 2012b; Li et al., 2016). Importantly, knowledge about these pathways could be harnessed to develop effective neuropeptide mimetics that might disrupt these pathways and serve as a new generation of biorational agents for pest control strategies.

The specificity of the ligand for its receptor arises from the recognition of the binding pocket of the GPCR. Unfortunately, only a few studies in insects have highlighted the structure-activity relationships of the receptor revealing features and residues that are vital for peptide ligand activation and specificity. Hence, to enhance our understanding of how the ACPR is selectively activated by its native ACP ligand, this study implements a structure-activity analysis through the generation of receptor chimera. Previously, a series of analogues based on the original ACP peptide of *A. aegypti* were designed and screened against the ACPR using a heterologous system to identify the crucial residues of the ligand necessary for receptor activation (Wahedi et al., 2019). The insight gained on the ACP ligand guided this research study that focuses instead on the ACP receptor (ACPR). Thus, in this current study, we used the mosquito *A. aegypti* as a model to study critical residues and features that facilitate specificity of ACPR for its ligand (ACP) while preventing promiscuous activation by the structurally- and evolutionarily-related peptide, AKH (Hansen et al., 2010; Hauser and Grimmelikhuijzen, 2014; Tian et al., 2016; Zandawala et al., 2018). The knowledge gained on the interaction between each ligand with their respective receptor may benefit development of selective biorational insecticides (Hill et al., 2018). The characterization of ACP and AKH neuropeptides could be utilized to design insecticides and pest control agents with improved specificity. Therefore, the main goal of this study was to determine how the ACP receptor (ACPR) elicits strong specificity for its native ligand in the *A. aegypti* mosquito by mutagenizing specific regions of the ACP receptor to identify features most critical for ligand-mediated activation and specificity. To achieve this, we first generated ACP receptor chimera using mutagenesis by singly replacing the three ACPR extracellular loops (ECL1, ECL2, and ECL3) in their entirety and then incorporating the corresponding ECL from the *A. aegypti* AKH receptor (AKHR) (Oryan et al., 2018). These receptor chimeras were then tested using a heterologous functional assay to confirm whether ACP was still capable of receptor activation after swapping the full extracellular loops. Secondly, following bioinformatic analysis involving multiple sequence alignment of various insect ACP and AKH receptors, we predicted specific residues in the ACPR extracellular loops that might be most critical for ligand fidelity and specificity in the mosquito *A. aegypti*. Three independent ACPR chimeras were generated after identifying highly-conserved residues within the three extracellular loops of insect ACP receptors and replacing them with residues highly conserved in insect AKH receptors. Thus, this study set out to test the following hypotheses: first, modification of extracellular loops of the ACPR determines binding sensitivity to its native ligand, ACP, while altering its sensitivity to other structurally-related ligands, such as AKH; second, ACPR chimera having highly-conserved residues within their extracellular loop replaced with corresponding highly-conserved residues from insect AKH receptors will exhibit reduced sensitivity to ACP while improving responsiveness to AKH.

## Materials and Methods

### Heterologous functional receptor assay

#### 1. Generation of A. aegypti ACP receptor chimera by site-directed mutagenesis

To create ACP receptor (ACPR) chimera, the native *A. aegypti* ACPR sequence was used to incorporate the three extracellular loops of the *A. aegypti* AKH receptor (AKHR) based on previously published sequences (Oryan et al., 2018; Wahedi et al., 2019; Wahedi and Paluzzi, 2018). The open reading frame (ORF) of the cloned *A. aegypti* ACPR was inserted into pBudCE4.1 mammalian expression vector following procedures described previously (Wahedi and Paluzzi, 2018; Wahedi et al., 2019). The ACPR chimeras were created by singly replacing the ACPR extracellular loops (ECLs) and incorporating ECLs of *Aedae*AKHR (see **Fig. 1**; schematic overview of the approach). The Consensus Constrained Topology Prediction (CCTOP) web server was used as a guide to predict positions of extracellular loops in order to design the forward and reverse primers used in PCR-based site-directed mutagenesis (Dobson et al., 2015). The forward and reverse primers were designed using the Primer3 module in Geneious® 6.1.8 Software (Biomatters Ltd, Auckland, New Zealand) (**Table 1**). The functional validation of the three ACPR chimeras was examined using a heterologous functional assay described previously (Wahedi and Paluzzi, 2018). Also, to identify the specific key residues that are highly conserved but unique between the ACP and the AKH receptors, which might be critical for ligand-binding specificity, the sequences of both these receptors in *A. aegypti* were compared with homologous sequences from other insects (see **Table 2**), with the analysis focusing on conserved residues within the extracellular loops (ECL1, ECL2, and ECL3) of these two distinct receptor families using the ClustalW online server. As mentioned above, the Primer3 module in Geneious® 6.1.8 Software was used to design primers (**Table 3**) to substitute only the highly conserved residues in order to generate the three selective ECLs chimeras of *Aedae*ACPR. Q5 High Fidelity DNA Polymerase (New England Biolabs, Whitby, ON) was used to amplify the three extracellular loop fragments and the native *Aedae*ACPR construct in pBudCE4.1 Gα15 was used as a template (Wahedi et al., 2019). PCR products (ACPR ECL1, ACPR ECL2 and ACPR ECL3) were assembled into pBudCE 4.1 Gα15 using NEBuilder HiFi DNA Assembly Cloning Kit (New England Biolabs, Whitby, ON) following the protocol guidelines. Briefly, each PCR product was mixed with HiFi DNA Assembly Master Mix and pBudCE 4.1 Gα15 (using a 2:1 insert:vector ratio) and then incubated in a thermocycler at 50°C for 30 minutes. Standard bacterial transformation was used with NEB 5-alpha competent *E. coli* cells (New England Biolabs, Whitby, ON) and grown on LB plates containing Zeocin™ antibiotic overnight at 37°C. The recombinant clones containing these constructs were then grown overnight in LB liquid media containing 25 μg/ml of Zeocin™ antibiotic and plasmid DNA isolation was isolated using the Monarch® Plasmid Miniprep Kit (New England Biolabs, Whitby, ON). Base-pair accuracy and orientation of the receptor constructs were confirmed by Sanger sequencing (Center for Applied, Genomics, Hospital for Sick Children, Toronto, ON). Endotoxin-free, high titer plasmid DNA was isolated using a ZymoPURE™ II Plasmid Midiprep Kit (Zymo Research, Irvine, USA) and used for transfection of mammalian cells utilized in the functional heterologous receptor assay.

**Figure 1.**
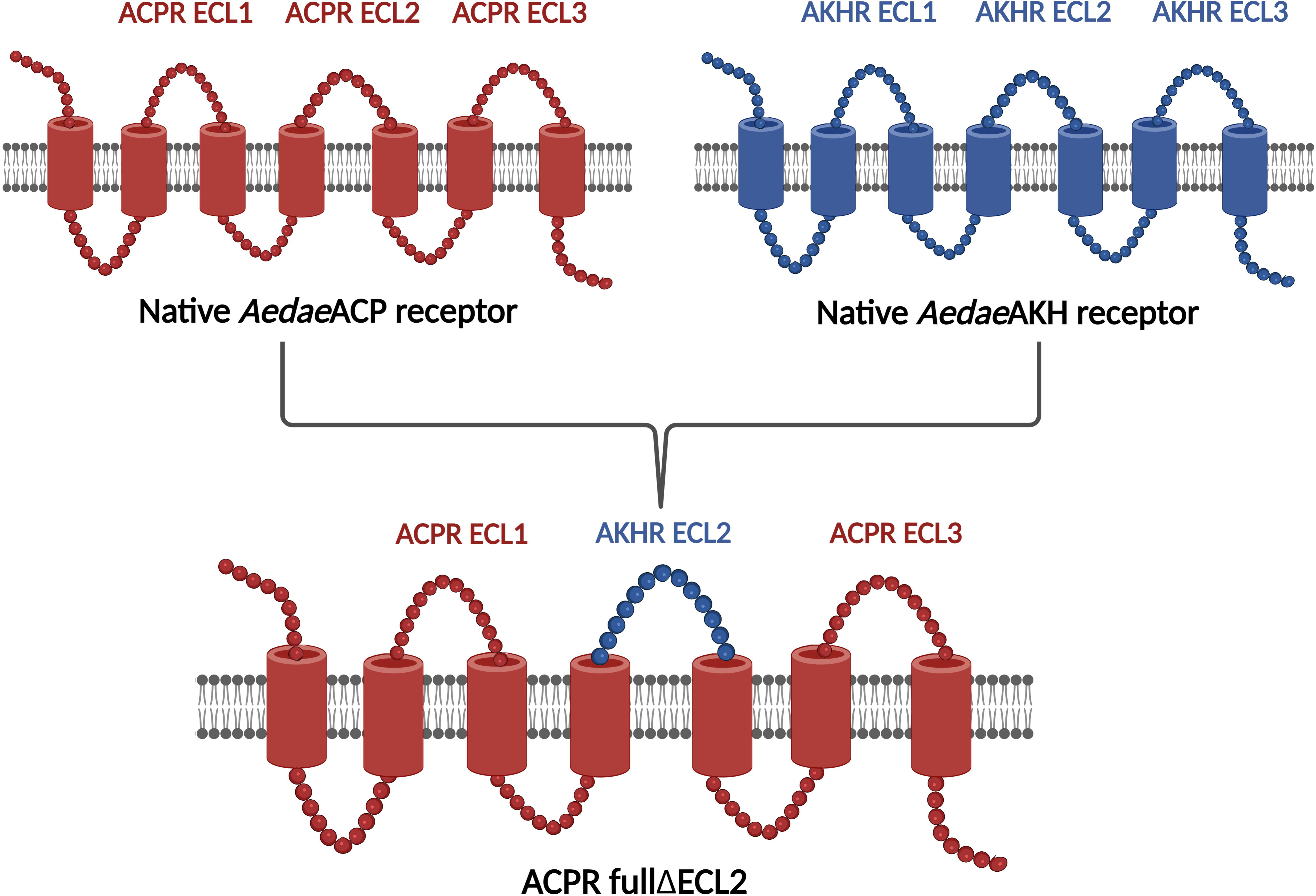
Schematic diagram showing design of *Aedae*ACPR chimera using the extracellular loop of *Aedae*AKHR and the native sequence of *Aedae*ACPR. The *Aedae*AKH receptor, including its three extracellular loops (AKHR ECL1, AKHR ECL2, AKHR ECL3) are presented in blue. The *Aedae*ACP receptor, including its three extracellular loops (ACPR ECL1, ACPR ECL2, and ACPR ECL3) are shown in red. For brevity, only the ACPR entire ECL2 chimera (ACPR fullΔECL2), with complete substitution of the second extracellular loop, is shown as an example at the bottom of the figure. Illustration was created using BioRender (BioRender.com).

**Table 1.**
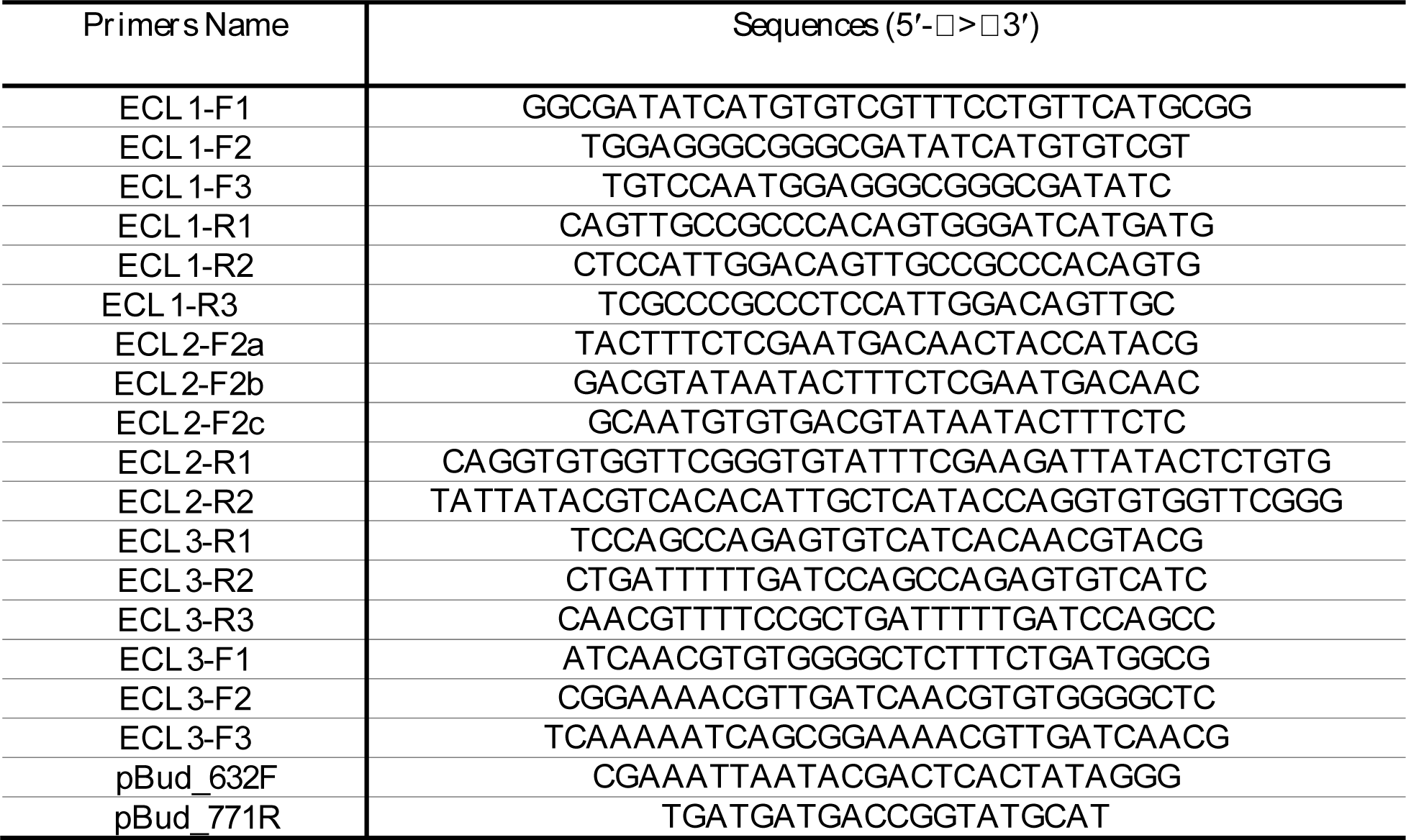
Primers used for mutagenesis creating the three ECLs chimera of *Aedae*ACPR.

**Table 2.**
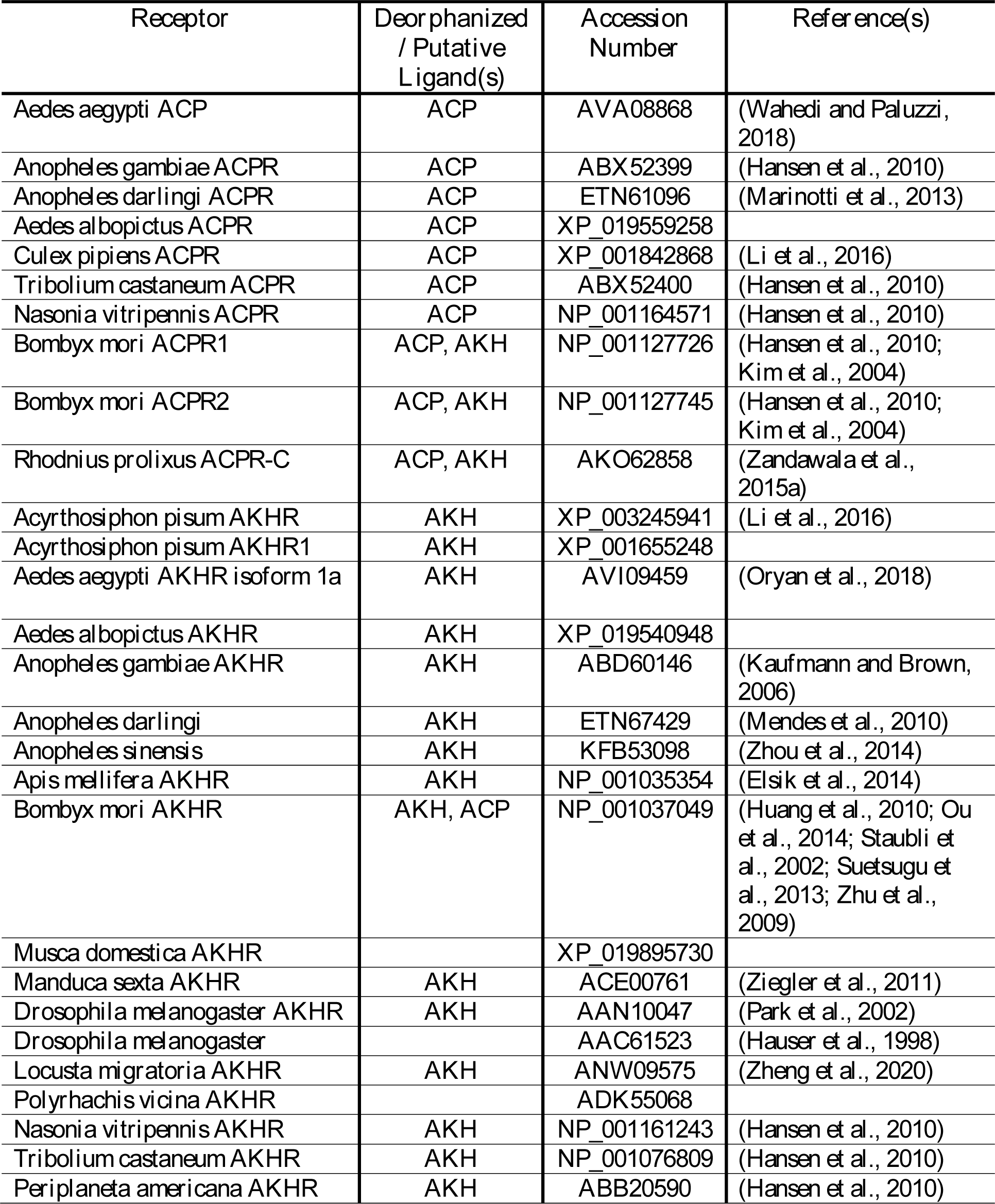
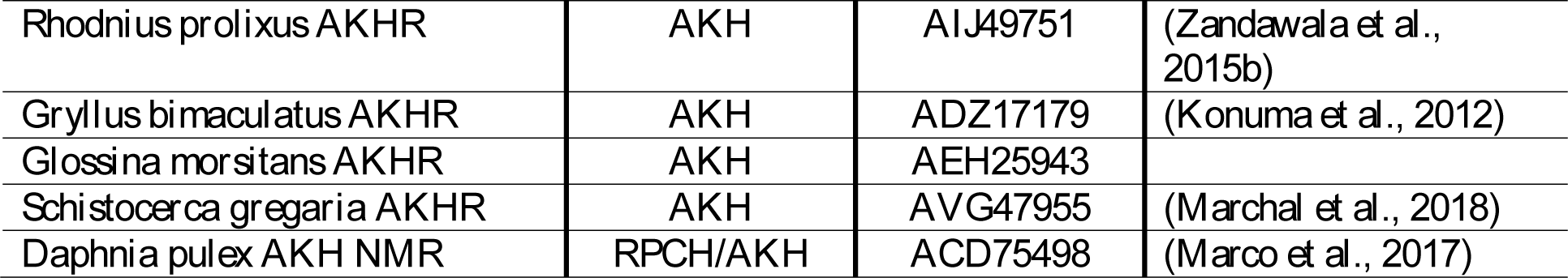
GenBank accession numbers and relevant literature references for the receptors used in the multiple sequence alignment (Fig. 3). The receptors that lack references were predicted by automated computational analysis.

| Receptor | Deorphanized / Putative Ligand(s) | Accession Number | Reference(s) |
| --- | --- | --- | --- |
| *Aedes aegypti* ACP | ACP | AVA08868 | (Wahedi and Paluzzi, 2018) |
| *Anopheles gambiae* ACPR | ACP | ABX52399 | (Hansen et al., 2010) |
| *Anopheles darlingi* ACPR | ACP | ETN61096 | (Marinotti et al., 2013) |
| *Aedes albopictus* ACPR | ACP | XP_019559258 |  |
| *Culex pipiens* ACPR | ACP | XP_001842868 | (Li et al., 2016) |
| *Tribolium castaneum* ACPR | ACP | ABX52400 | (Hansen et al., 2010) |
| *Nasonia vitripennis* ACPR | ACP | NP_001164571 | (Hansen et al., 2010) |
| *Bombyx mori* ACPR1 | ACP, AKH | NP_001127726 | (Hansen et al., 2010; Kim et al., 2004) |
| *Bombyx mori* ACPR2 | ACP, AKH | NP_001127745 | (Hansen et al., 2010; Kim et al., 2004) |
| *Rhodnius prolixus* ACPR-C | ACP, AKH | AKO62858 | (Zandawala et al., 2015a) |
| *Acyrtosiphon pisum* AKHR | AKH | XP_003245941 | (Li et al., 2016) |
| *Acyrtosiphon pisum* AKHR1 | AKH | XP_001655248 |  |
| *Aedes aegypti* AKHR isoform 1a | AKH | AVI09459 | (Oryan et al., 2018) |
| *Aedes albopictus* AKHR | AKH | XP_019540948 |  |
| *Anopheles gambiae* AKHR | AKH | ABD60146 | (Kaufmann and Brown, 2006) |
| *Anopheles darlingi* | AKH | ETN67429 | (Mendes et al., 2010) |
| *Anopheles sinensis* | AKH | KFB53098 | (Zhou et al., 2014) |
| *Apis mellifera* AKHR | AKH | NP_001035354 | (Elsik et al., 2014) |
| *Bombyx mori* AKHR | AKH, ACP | NP_001037049 | (Huang et al., 2010; Ou et al., 2014; Staubli et al., 2002; Suetsugu et al., 2013; Zhu et al., 2009) |
| *Musca domestica* AKHR |  | XP_019895730 |  |
| *Manduca sexta* AKHR | AKH | ACE00761 | (Ziegler et al., 2011) |
| *Drosophila melanogaster* AKHR | AKH | AAN10047 | (Park et al., 2002) |
| *Drosophila melanogaster* |  | AAC61523 | (Hauser et al., 1998) |
| *Locusta migratoria* AKHR | AKH | ANW09575 | (Zheng et al., 2020) |
| *Polyrhachis vicina* AKHR |  | ADK55068 |  |
| *Nasonia vitripennis* AKHR | AKH | NP_001161243 | (Hansen et al., 2010) |
| *Tribolium castaneum* AKHR | AKH | NP_001076809 | (Hansen et al., 2010) |
| *Periplaneta americana* AKHR | AKH | ABB20590 | (Hansen et al., 2010) |
| *Rhodnius prolixus* AKHR | AKH | AIJ49751 | (Zandawala et al., 2015b) |
| *Gryllus bimaculatus* AKHR | AKH | ADZ17179 | (Konuma et al., 2012) |
| *Glossina morsitans* AKHR | AKH | AEH25943 |  |
| *Schistocerca gregaria* AKHR | AKH | AVG47955 | (Marchal et al., 2018) |
| *Daphnia pulex* AKH NMR | RPCH/AKH | ACD75498 | (Marco et al., 2017) |

**Table 3.**
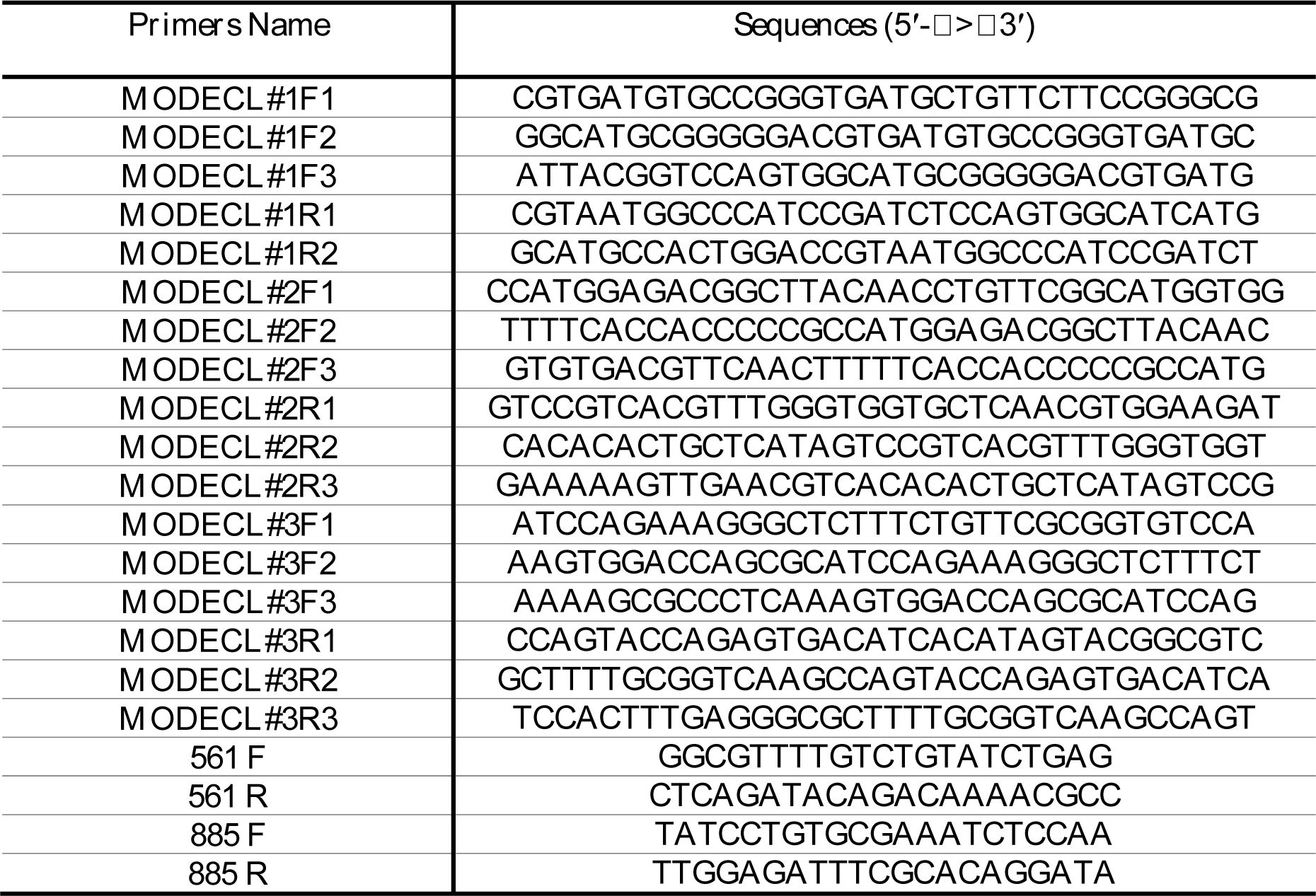
Primers used for site-directed mutagenesis to substitute specific residues of ACPR ECLs that are highly conserved and may be important for ligand specificity of *Aedae*ACPR.

#### 2. Cell culture, transfections, and calcium bioluminescence receptor assay

Functional activation of the three complete ECL *Aedae*ACP receptor chimeras (ACPR ECL1, ACPR ECL2 and ACPR ECL3), as well as the three selectively modified *Aedae*ACPR ECLs chimeras, were carried out following a previously described protocol utilizing a Chinese hamster ovary (CHO)-K1 cell line stably expressing aequorin (Paluzzi et al., 2012). Cells were grown in Dulbecco’s modified eagles medium: nutrient F12 (DMEM: F12) media containing 10% heat-inactivated fetal bovine serum (FBS; Wisent, St. Bruno, QC), 200μg/mL geneticin, and antimycotic-antibiotic mixture as reported previously (Wahedi and Paluzzi, 2018). Cells were grown to 90% confluency and were transfected with the native ACPR, the full ECL ACPR chimera (ECL1-3) and the selectively modified ACPR ECLs chimera using Lipofectamine 3000 transfection reagent (Invitrogen, Burlington, ON) using a 1:3 DNA (μg) to transfection reagent volume (μL) ratio. A pcDNA mCherry construct was used as a control to assess transfection efficiency and to further validate activation specificity of the ACPR ECLs chimeras. Cells were harvested for the functional assay by detaching from the culture flasks at 48 hours post-transfection using 5mM ethylenediaminetetraacetic acid (EDTA) in Dulbecco’s PBS. Cells were prepared for the receptor functional assay as described previously (Oryan et al., 2018; Wahedi and Paluzzi, 2018).

Cells prepared for the functional assay were loaded into each well of a white 96-well plate using an automated injector unit, and the luminescent response was measured with a Synergy 2 Multi-Mode Microplate Reader (BioTek, Winooski, VT, USA). The *Aedae*ACP (pQVTFSRDWNAa) and *Aedae*AKH (pQLTFTPSWa) peptides were commercially synthesized (purity >90%; Genscript, Piscataway, NJ) and diluted in BSA assay media (0.1% BSA in DMEM: F12) to achieve the final working concentration. Luminescent response following application of 1μM *Aedae*ACP or *Aedae*AKH peptides onto the native ACPR, each of the three full ECL substituted *A. aegypti* ACPR chimera as well as the selectively substituted *A. aegypti* ACPR chimera were examined. Negative controls were carried out using BSA assay media alone, whereas 50μM ATP prepared in BSA assay media served as a positive control, which acts on endogenous purinoceptors, as previously described (Lajevardi and Paluzzi, 2020). Background luminescence was subtracted from luminescent responses following peptide challenge, which was then normalized relative to the maximal luminescent response obtained using ATP.

### Statistical analyses

Data was compiled using Microsoft Excel and transferred to GraphPad Prism 8.02 (GraphPad Software, San Diego, USA) to create all figures and conduct statistical analyses. Receptor functional assay average luminescence data were analyzed by performing either a t-test or one-way ANOVA followed by Tukey’s multiple comparison post-test to compare control and different experimental treatments. In all statistical tests, p<0.05 was considered significant.

## Results

### Ligand-receptor interaction heterologous functional assay

#### 1. Independent substitution of the three ECLs of AedaeAKH receptor into the ACPR chimera

The first step for determining the ACP receptor (ACPR) specificity and understanding the particular regions of the ACPR most critical for ligand fidelity and specificity was creating ACPR chimera incorporating extracellular loops from the *Aedae*AKH receptor. The functional assay aimed to confirm whether ACP was still functional and capable of activating the ACPR chimera after individually swapping out the complete ACPR ECL1, 2 and 3 with the corresponding ECLs of the AKH receptor. Using a heterologous functional assay, the luminescent response of CHO-K1 aequorin cells expressing the native *A. aegypti* ACPR or one of the three whole ECL ACPR chimera, namely ACPR fullΔECL1, ACPR fullΔECL2 and ACPR fullΔECL3), was used to validate their activation. Nucleotide and deduced amino acid sequences of these three chimera are available in supplementary data files. The results confirmed that the ACPR elicited the highest response to *Aedae*ACP (p < 0.0001), as this peptide binds with its native receptor (**Fig. 2A**). In contrast, the three full ECL ACPR chimera showed no response with either the *Aedae*ACP (**Fig. 2A**) or *Aedae*AKH (**Fig. 2B**).

**Figure 2.**
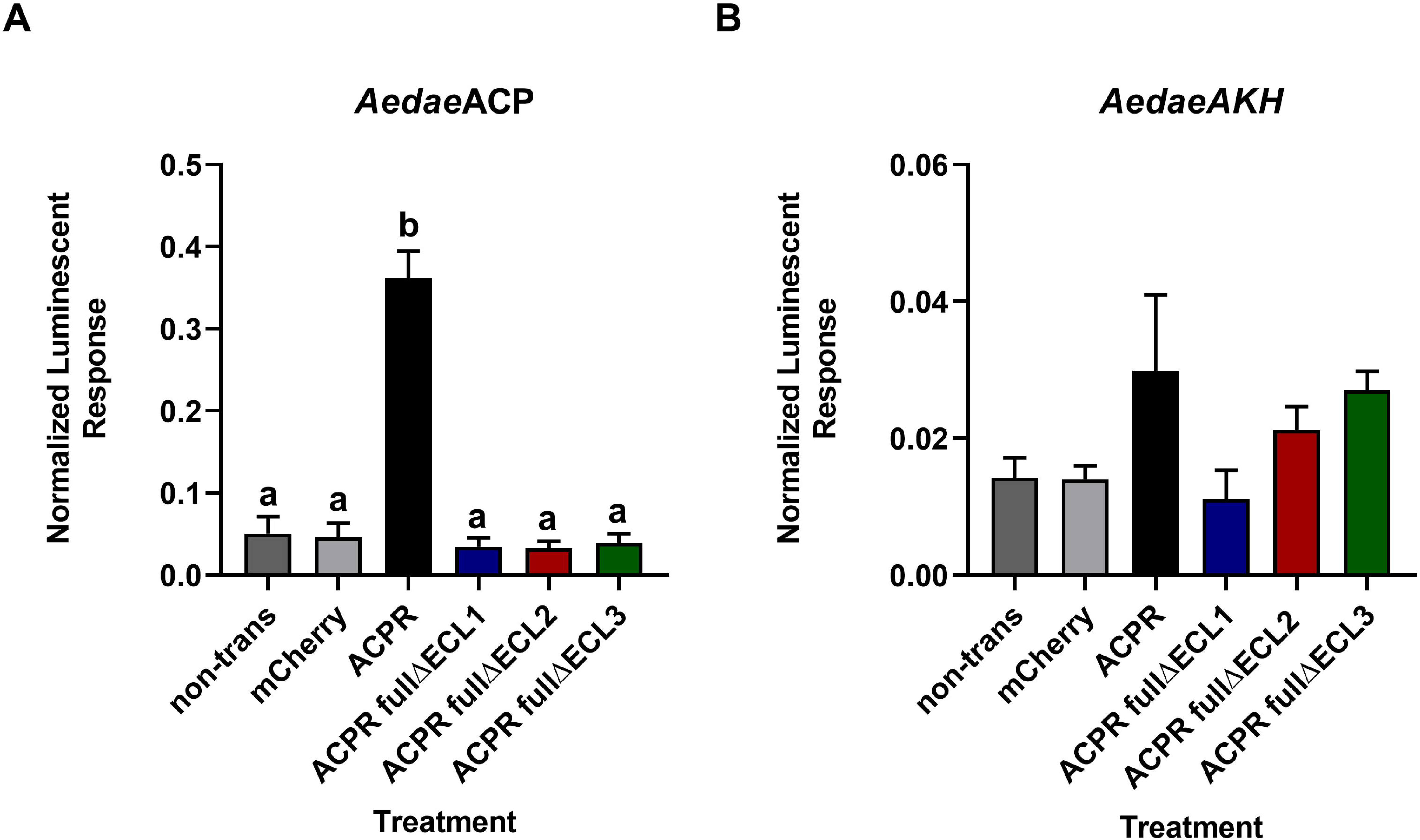
Functional heterologous receptor assay expressing either the native *A. aegypti* ACPR or one of three full ECL chimeras of the ACPR. (A) Normalized luminescent response of the native ACPR and three ACPR ECLs chimera after treatment with 1μM *Aedae*ACP. **(B)** Normalized luminescent response of the native ACPR and the three ACPR ECL chimera after treatment with 1μM *Aedae*AKH. Treatments included non-transfected control (non-trans), transfected control (mCherry), native ACP receptor (ACPR), ACPR ECL1 chimera (ACPR fullΔECL1), ACPR ECL2 chimera (ACPR fullΔECL2), and ACPR ECL3 chimera (ACPR fullΔECL3). Background luminescent response (BSA control) was subtracted from experimental samples and normalized relative to the maximal luminescence following treatment with ATP (10^−5^ M). Different letters denote bars that are significantly different from one another, as determined by a one-way ANOVA and Tukey’s multiple comparison post-test (p < 0.05). Data represent the mean ± standard error (n=3).

#### 2. Identification of specific key residues in the three ECLs of ACPR

Considering the dramatic impact on receptor activation by ACP following the individual substitution of the full individual extracellular loop domains, we next implemented a more targeted approach. Specifically, we aimed to identify and substitute specific highly-conserved residues in each of the three ECLs rather than replacing the full extracellular loop in their entirety. We reasoned this might help to identify key residues in the three ECLs that may confer ligand binding specificity. Thus, we aimed at discerning specific critical residues necessary for functional ligand-binding that are highly conserved and important for ligand specificity. Multiple sequence alignment of ACP and AKH receptors from various insect species revealed putative residues within the three ECLs that may contribute towards ligand specificity (**Fig. 3**). For instance, the valine residue (V_165_ in *A. aegypti* ACPR; Wahedi and Paluzzi, 2018) that is highly conserved in this position of the ECL1 in insect ACP receptors was substituted with an isoleucine (I_131_ in both *A. aegypti* AKHR biologically active isoforms, AKHR-IA and AKHR-II; Oryan et al., 2018), which is highly conserved in AKH receptors across insects **(Fig. 3A)**; and similar substitutions of highly conserved residues were performed along the entire region of the three ECLs. GenBank accession numbers and related references for insect ACPR or AKHR sequences used for the multiple sequence alignment are listed in **Table 2**.

**Figure 3.**
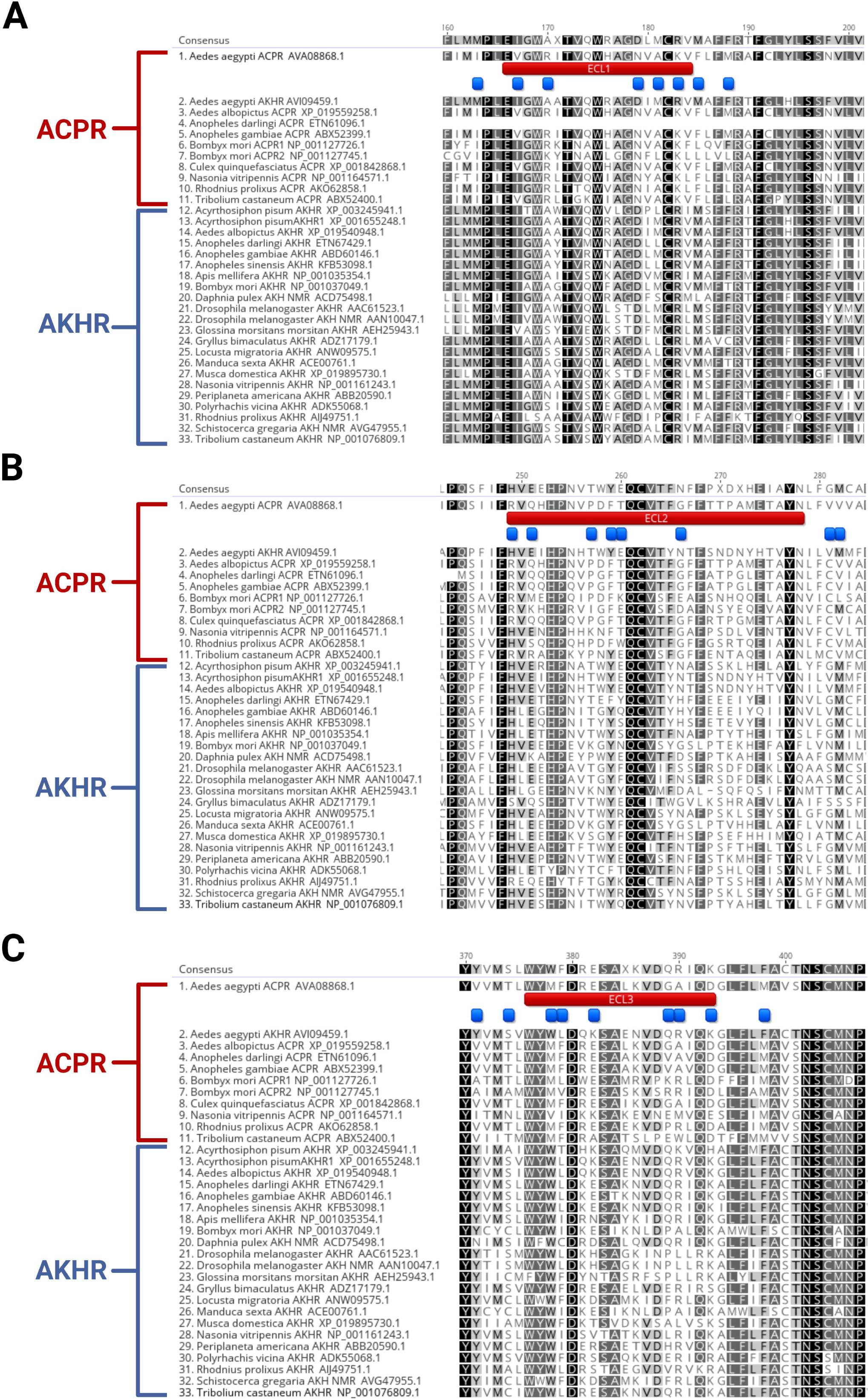
Sequence alignment of select insect ACP receptors (ACPR) and AKH receptors (AKHR) to identify specific residues that are highly conserved and could be important for the ligand specificity conferred by the extracellular loops. The three extracellular loop regions, **(A)** ECL1, **(B)** ECL2 and **(C)** ECL3, with residues outlined on the top with blue boxes indicating receptor-specific residues which are highly conserved. Highlighting of residues indicates % identity with black denoting 100% sequence identity, dark grey denotes 80 > 100% identity, light grey represents amino acid positions with 60 > 80% sequence identity, and white denotes amino acid positions with < 60% sequence identity. Receptor protein sequences are labelled by species name and identified with their GenBank accession number. References for sequences are listed in Table 2.

Subsequently, three new independent ACPR-ECL chimeras were designed to modify these specific residues in the three ECLs. Nucleotide and deduced amino acid sequences of the various chimera are available in supplementary data files. Interestingly, the ACPRΔECL1 selective chimera showed the same response as the native ACPR to *Aedae*ACP (**Fig. 4A)**, while the selective ACPRΔECL2 chimera showed a significantly higher response (p < 0.0001) to *Aedae*ACP compared to the native ACPR (**Fig. 4A)**. In contrast, the selective ACPRΔECL3 chimera showed a significantly decreased response (p < 0.0001) to *Aedae*ACP compared to the response by the native ACPR (**Fig. 4A**). Notably, all three of the selectively modified ACPR chimera showed a significantly higher response to *Aedae*AKH than the native ACPR response. Specifically, the ACPRΔECL2 and ACPRΔECL3 chimeras demonstrated the highest response (p <0.0001) to *Aedae*AKH, being also significantly higher than the response observed with ACPRΔECL1 (p = 0.0348) (**Fig, 4B**).

**Figure 4.**
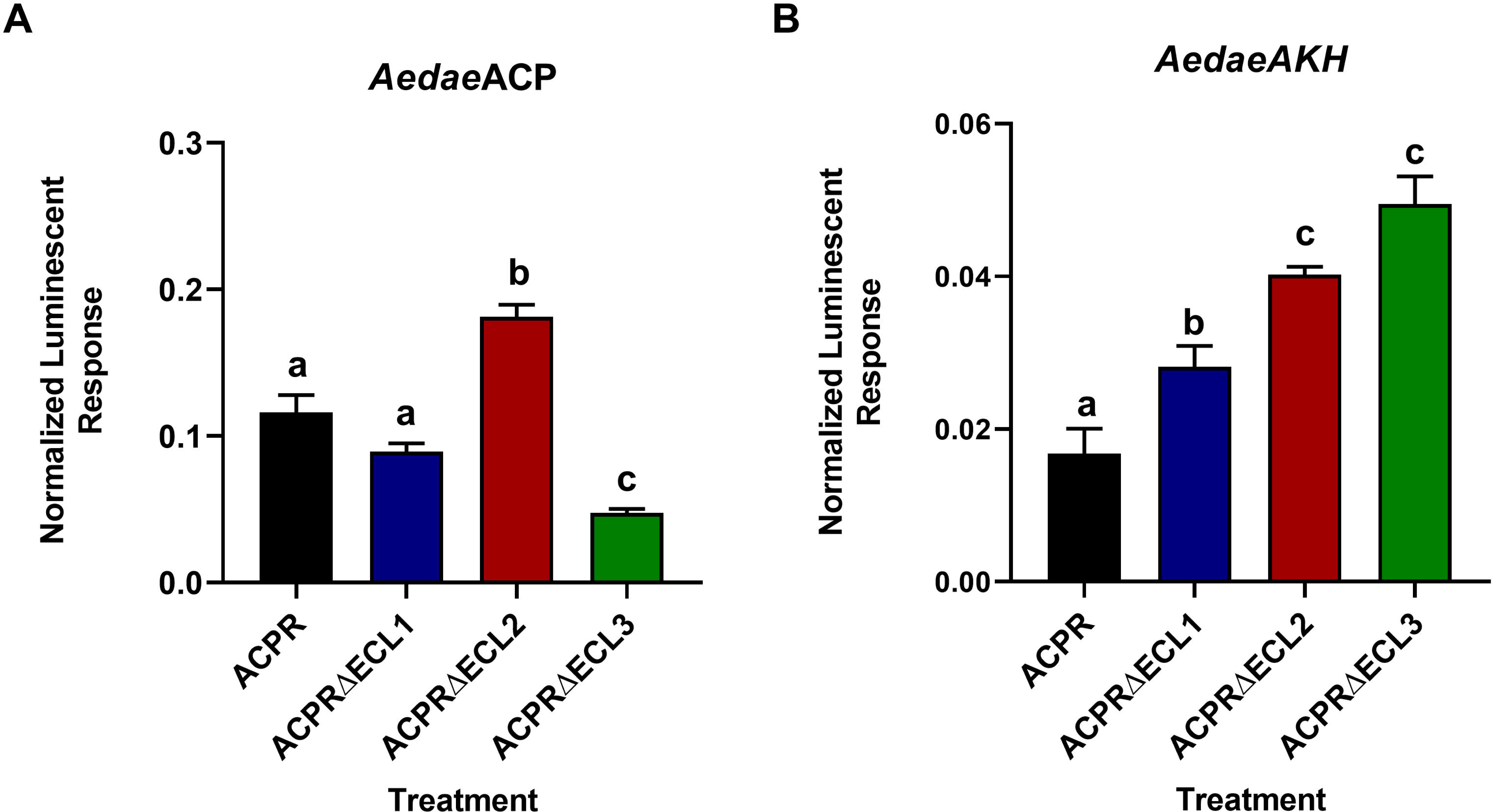
Bioluminescent responses of CHO-K1 aequorin cells transiently expressing either the native *A. aegypti* ACPR or the three chimeras with selectively modified residues within the ECLs. (A) Normalized luminescence of the three independent ECLs chimera following treatment with 1μM *Aedae*ACP. **(B)** Normalized luminescence of the three independent ECLs chimera after treatment with 1μM *Aedae*AKH. Background luminescence (i.e. BSA control) was subtracted from experimental samples and data was then normalized relative to the maximal luminescent response following ATP treatment (10^−5^ M). Different letters denote bars that are significantly different from one another as determined by a one-way ANOVA and Tukey’s multiple comparison post-test (p < 0.05). Data represent the mean ± standard error (n=3).

#### 3. Combination of two selectively modified ECLs of the ACPR chimeras

After observing the activity of ACPR chimera with selectively modified residues in individual ECLs, the next aim was to determine the effect of the combination of two selectively modified ACPR ECL chimeras. This involved a combination of ACPR ECL1 chimera with ACPR ECL2 chimera (ACPRΔECL1+2), ACPR ECL2 chimera with ACPR ECL3 chimera (ACPRΔECL2+3) and lastly, ACPR ECL1 chimera with ACPR ECL3 chimera (ACPRΔECL1+3). The results demonstrated that all double ECL chimera showed a significantly decreased response to *Aedae*ACP compared to the native ACPR response (p < 0.0001) (**Fig. 5A**). On the other hand, the ACPRΔECL2+3 or ACPRΔECL1+3 double chimera showed the same response to *Aedae*AKH as the native ACPR response. Notably, however, the ACPRΔECL1+2 double chimera showed a significantly higher response (p = 0.0407) to *Aedae*AKH compared to the native ACPR (**Fig. 5B**).

**Figure 5.**
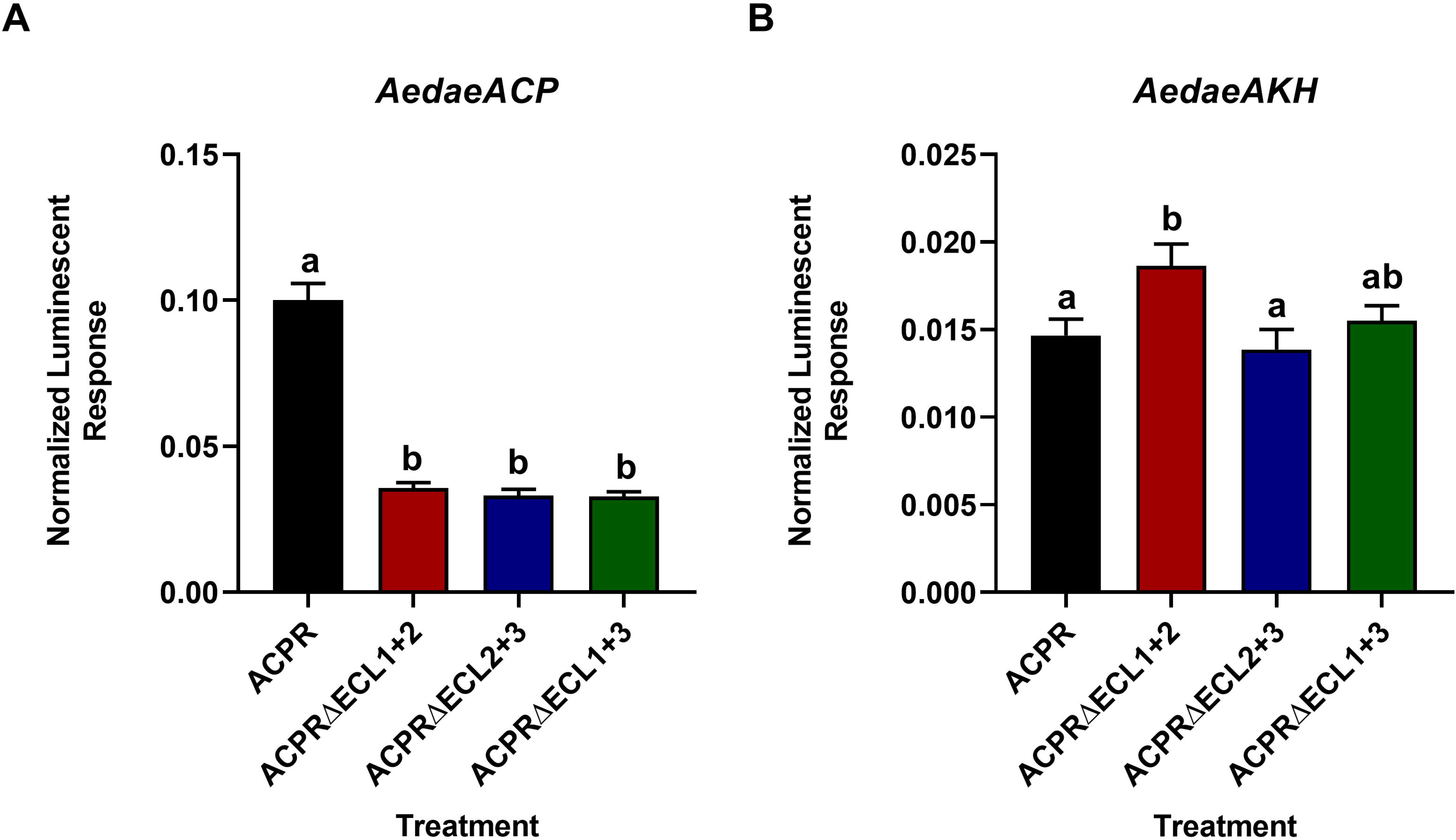
Receptor functional assay of CHO-K1 aequorin cells transiently expressing either the native *A. aegypti* ACPR or the combination of two selectively modified ECLs of the ACPR chimeras. **(A)** Normalized luminescent response of the ACPR chimera combining two selectively modified ECLs following treatment with 1μM *Aedae*ACP. **(B)** Normalized luminescent response of the ACPR chimera combining two selectively modified ECLs following treatment with 1μM *Aedae*AKH. Background luminescent response obtained following the vehicle control (BSA media) was subtracted from the experimental samples and data was then normalized relative to the maximal luminescence following ATP treatment (10^−5^ M). Different letters denote bars that are significantly different from one another as determined by a one-way ANOVA and Tukey’s multiple comparison post-test (p < 0.05). Data represent the mean ± standard error (n=3).

#### 4. Combination of all three selectively modified ECLs of the ACPR chimera

A combination of the three selectively modified ACPR ECLs chimeras (ACPRΔECL1+2+3) was tested to determine the effect of these combined specific residue substitutions on receptor sensitivity and activation by the two closely-related ligands, *Aedae*ACP and AKH. The results revealed a significantly decreased response for the combined chimera ACPRΔECL1+2+3 compared to the native ACPR response to *Aedae*ACP (p < 0.0001) (**Fig. 6A**). Comparatively, the combined chimera ACPRΔECL1+2+3 resulted in a significantly increased response to *Aedae*AKH (p = 0.0217) (**Fig. 6B**).

**Figure 6.**
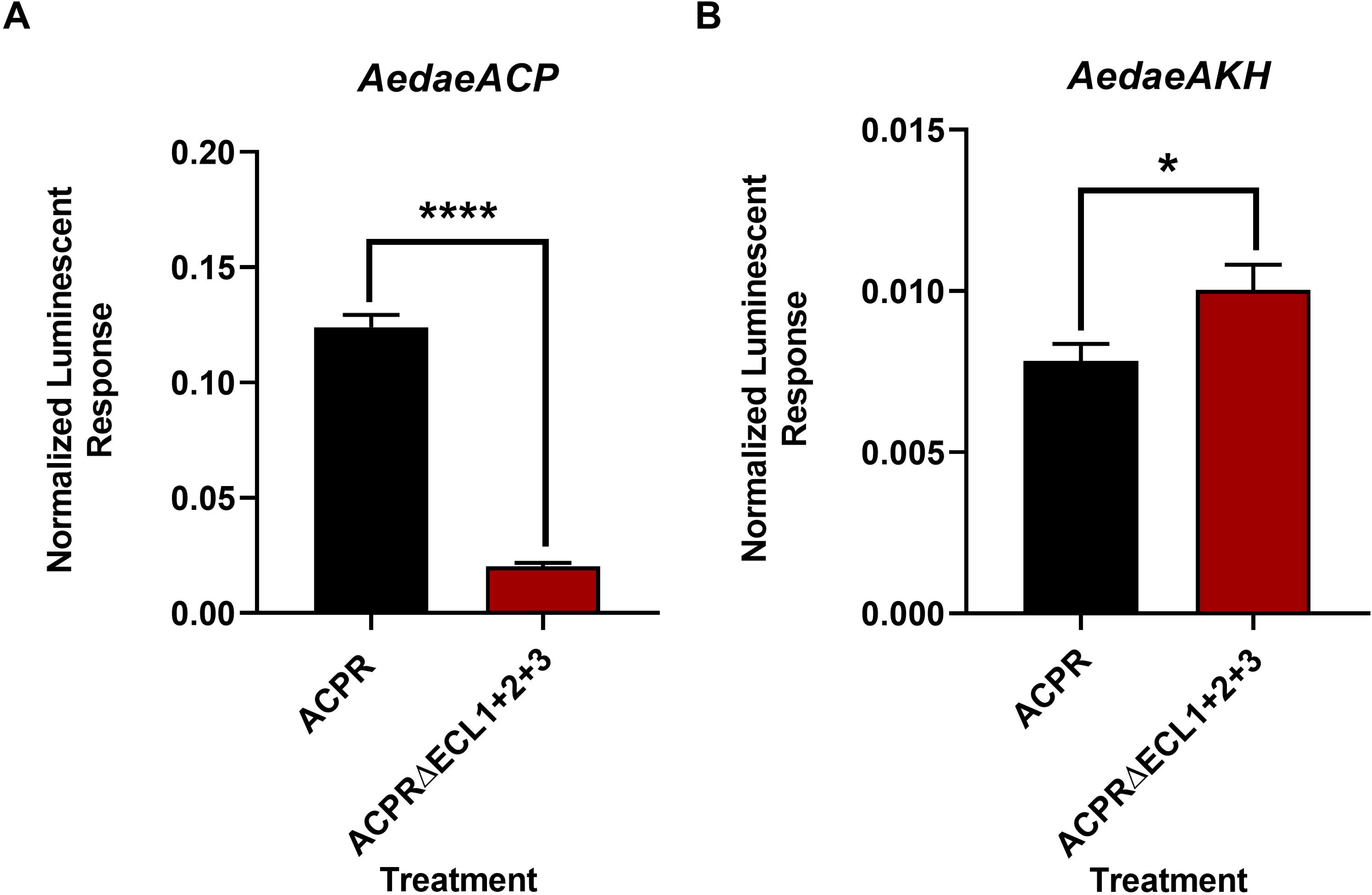
Bioluminescent responses of CHO-K1 aequorin cells transiently expressing either the native *A. aegypti* ACPR or variant combining all three selectively modified ECLs of the ACPR chimeras. (A) Normalized luminescent response of the ACPR chimera combining all three selectively modified ECLs following treatment with 1μM *Aedae*ACP. **(B)** Normalized luminescent response of the ACPR chimera combining all three selectively modified ECLs following treatment with 1μM *Aedae*AKH. Background luminescent response (BSA control) was subtracted from experimental samples and data was normalized relative to the maximal luminescent response following treatment of ATP (10^−5^ M). Data represent meanL±Lstandard error (n=3), (*) denotes significance (p<0.05) as determined using an unpaired two-tailed t-test.

## Discussion

The adipokinetic hormone/corazonin-related peptide (ACP) is structurally intermediate between AKH and CRZ (Hansen et al., 2010). Previous *in vitro* studies characterizing AKH receptor (AKHR) in the silkworm *B. mori* have demonstrated that high concentrations of AKH can activate the ACP receptor (ACPR) and *vice versa* (Zhu et al., 2009; Shi et al., 2011), which indicates that ACP and AKH are closely related to each other (Hansen et al., 2010). In arthropods, despite ACP and AKH being part of two distinct signaling systems, there is a notable similarity between these two peptides (Hansen et al., 2010). This similarity caused ACP to be misclassified previously as a member of the AKH peptide family in several studies prior to the discovery of the ACP receptor (Siegert, 1999; Kaufmann and Brown, 2006; Kaufmann et al., 2009). Moreover, extensive structure-activity studies were carried out on AKH receptors in several species using natural and synthetic analogs of the AKH neuropeptide family (Ziegler et al., 1991, 1998; Gäde, 1992; Poulos et al., 1994; Lee and Goldsworthy, 1996; Lee et al., 1997; Gäde et al., 2000; Velentza et al., 2000; Caers et al., 2012a; Marco and Gäde, 2015, 2019). On the other hand, limited data on biological actions or structure-activity relationships is available for the ACP system in insects.

A recent study examined the structure-activity relationship (SAR) of the ACP ligand in *A. aegypti*, using the ACP analogs as well as the natural AKH isoforms as ligands, to determine the ligand characteristics necessary for ACP receptor activation and specificity (Wahedi et al., 2019). However, the SAR of the ACP receptor in *A. aegypti* has not yet been investigated. Consequently, the present study intended to advance our understanding of SAR of two GnRH-related systems in *A. aegypti* sharing the most recent evolutionary origin that sustain independence of function and signaling despite their relatively high degree of ligand and receptor homology (Hansen et al., 2010; Hauser and Grimmelikhuijzen, 2014; Tian et al., 2016; Zandawala et al., 2018). Specifically, this study set out to examine the SAR of the ACPR in the *A. aegypti* mosquito by substitution of key regions or selected residues of the receptor. In doing so, this provides insight into the specific structural features necessary for ligand fidelity and sustaining independence of the ACP and AKH signaling systems in insects.

As a consequence of the sequence conservation between the ACP and AKH signaling system, specific regions of the ACPR potentially most critical for ligand specificity were examined by singly replacing the three extracellular loops (ECLs) of the native ACPR and incorporating those from the AKHR of *A. aegypti*. Overall, it was observed that the three ACPR whole ECL chimera receptors showed no response to either AKH or ACP, indicating these ligands could no longer bind or the chimera were non-functional when a single full extracellular loop of ACPR (ECL 1, 2 and 3) was switched with that from the AKHR. This finding suggests that the complete replacement of each ECL has a detrimental effect on functional ligand-receptor binding. Thus, these results reveal that all three extracellular domains play a critical role in forming the ACP ligand-binding pocket. In agreement with these results, it is well documented for a variety of G-protein coupled receptors (GPCRs) that the ECLs play a key role in the activation and ligand-binding recognition (Wheatley et al., 2012). For example, one study focused on how the modifications of ECLs impacted ligand-binding and activation of GnRH by substituting the ECL2 domain of the human GnRH receptor with that of the chicken establishing the chicken GnRH ECL2 domain is responsible for conferring agonist activity to mammalian GnRH antagonists (Sun et al., 2001).

As a subsequent step following the replacement of the whole individual ECLs, highly conserved residues within the three ECLs of ACPR and AKHR in several insects were identified, and then modified to generate three new independent ACPR ECLs chimeras with these selected substitutions. Thus, the most highly conserved residues between ACP and AKH receptors were targeted in order to discriminate particular residues critical for ligand-binding and functionality. The results suggest that selectively modified residues within the third extracellular loop in particular appear to be the most critical for ACP activation of its receptor. This is not surprising since the ECL3 of GPCRs is more than just a connector between two transmembranes, but it is also involved in receptor activation and ligand selectivity (Lawson and Wheatley, 2004). Moreover, another study observed a similar finding to the current results but in another insect neuropeptide system using chimera receptors to determine the function of the extracellular loops in the pheromone biosynthesis-activating neuropeptide (PBAN)/pyrokinin GPCRs from insects (Choi et al., 2007). PBAN is a peptide considered to be a pyrokinin-2 (PK2) family member that is known to stimulate hindgut motility and induce the biosynthesis of sex pheromone in many insects; but different from the diapause hormone (DH), which is a pyrokinin-1 (PK1) family member (Raina et al., 1989; Rafaeli and Jurenka, 2003; Zhang et al., 2004a, 2004b; Choi et al., 2007; Jurenka, 2015; Ahn and Choi, 2018; Lajevardi and Paluzzi, 2020). After generating chimera receptors by swapping the third extracellular domain between a PBAN-receptor from a moth and pyrokinin-receptors from *Drosophila*, similar observations to the present study were reported indicating the third extracellular loop of these receptors is essential for recognition of the native peptide ligand (Choi et al., 2007). Separately, a more recent study revealed that the replacement of key proline residue (Pro_238_) located in the third ECL in *D. melanogaster* sex peptide receptor (*Drm*SPR) with the corresponding leucine residue (Leu_310_) from mosquitoes *A. aegypti* sex peptide receptor *(Aea*SPR) reduced its sensitivity to the sex peptide (SP) without changing its sensitivity to the ancestral ligand, the myoinhibitory peptide (MIP) (Lee et al., 2020). Therefore, further research will be needed to identify the specific amino acid residues that are the most critical in the third ECL of ACPR necessary for ligand binding and activation in *A. aegypti*.

A recent *in silico* study done on the stick insect, *Carausius morosus*, using the AKH receptor model, showed that crucial residues in ligand binding were located within the second ECL and the sixth and seventh transmembrane domains (Birgul Iyison et al., 2020). Using molecular modeling and mutagenesis studies, a study involving the human GnRH receptor suggested that the junction between the fourth transmembrane domain and second ECL plays a critical role in the peptide ligand binding and also in the conformational selection of the receptor (Forfar and Lu, 2011). Consequently, we decided to examine the effect of combining two ACPR ECL chimeras. Interestingly, the results revealed that all doubly combined ACPR ECLs chimeras showed a significantly decreased response to *Aedae*ACP compared to the native ACPR response. On the other hand, with regards to the *Aedae*AKH response, a few observations can be highlighted. First, ACPR chimeras combining selected substitutions to either ECL2 and ECL3 or ECL1 and ECL3 showed the same response as the ACPR, while the combination of the ACPR ECL1 and ECL2 showed a significantly increased response to *Aedae*AKH compared to the native ACPR response. Thus, the interactions between the first two selectively modified ECLs may play a key role in enhancing the response of ACPR to the AKH ligand. Lastly, a combination of the three ACPR ECLs chimeras demonstrated a significantly reduced response to *Aedae*ACP, while in contrast, a significantly increased response to *Aedae*AKH. Therefore, this study has identified several key residues of ACPR that are fundamental for ACP ligand binding fidelity while simultaneously elucidating the residues which help to prevent promiscuous binding by the closely related peptide, AKH. Consequently, findings from our research demonstrate that modifying ECLs of the ACP receptor to more closely resemble the ECLs from the AKH receptor leads to reduced sensitivity to binding of its native ligand, ACP, while improving responsiveness to AKH. Therefore, these data are highly important for studies examining the potential use of ACPR or AKHR as targets for developing novel biorational compounds, such as peptidomimetics, useful in pest control strategies.

### Concluding remarks and future directions

Taken together, until this present study, no structure-activity relationship analyses had been carried out on the ACP receptor in any arthropod. While not all the specific residues substituted within the extracellular loop domains may be directly involved in peptide binding, they might contribute towards maintaining the appropriate conformational structure that allows the binding of the peptide with its receptor. This current study emphasized that all three extracellular loop domains of the *A. aegypti* ACPR are indeed necessary to facilitate binding with its corresponding ligand. Notably, residues within the third extracellular loop domain appear to be the most critical for the ACP ligand-binding and recognition. Furthermore, a highly detrimental effect in the ACP binding recognition was observed after the combination of the three ACPR ECLs chimeras, while contrarily, an increased response occurred toward *Aedae*AKH. Consequently, this study succeeded in identifying several residues within ACPR that are pivotal for the ACP ligand-induced activation of this receptor. Collectively, the data obtained in this current study set the framework toward understanding the ACP ligand-binding pocket and also provide fundamental insight regarding the residues that play a key role in the activation of the ACP and AKH receptors by their corresponding ligands in *A. aegypti* mosquitoes. However, which of these specific amino acids are most important for ACP ligand binding within each extracellular domain of ACPR remains to be elucidated. Specific amino acid residues in ECLs could play a crucial role in agonist and/or antagonist binding to GPCRs (Hjorth et al., 1994). Exploring the indispensable amino acid residues of these two GnRH-related receptors in *A. aegypti* mosquito that are fundamental for activation by their respective ligands would help to clarify how these two evolutionarily related systems uphold specific signaling networks avoiding their cross-activation onto the structurally related but functionally distinct signaling systems. Hence, future investigations could focus on determining specific amino acids in each extracellular loop individually, with a specific focus on the third ECL that was determined herein to be the most critical. Thus, point mutations or site-directed mutagenesis of a subset of the highly conserved residues within each extracellular domain will be required to provide further insight for modeling the ligand-receptor structural interaction. This could confirm the most indispensable residues that are directly involved in the recognition and binding to the ACP ligand. Moreover, the next steps for future research could include utilizing nuclear magnetic resonance (NMR), as was used for the orthologs of AKH and its receptor in other species (Jackson et al., 2019, 2018; Jackson and Gäde, 2021). Thus, an approach involving molecular modeling could help determine the structural similarities and differences between *A. aegypti* ACPR and AKHR, which might shed light on the molecular interaction explaining how these receptors bind with their unique ligands. Moreover, it might give insight into design of some high-affinity agonists or antagonists to block the function of ACP and AKH, which could further facilitate the future design of insecticides against the same receptor of several insects.

## Acknowledgments

This research was supported by a Natural Sciences and Engineering Research Council of Canada (NSERC) Discovery Grant to JPP (RGPIN-2020-06130).

## Data availability

All data generated or analysed during this study are included in this published article (and its Supplementary Information files).

