## Supplementary information for "Insights into adipokinetic hormone/corazonin-related peptide receptor specificity and key residues for its activation in the human disease vector *Aedes aegypti* mosquito"

**ECL1- highlighted with Yellow color.**

**ECL2- highlighted with Bright Green color.**

**ECL3- highlighted with Turquoise color.**

**Selectively modified residues are denoted with red colored font.**

**Native *A. aegypti* ACPR** *(***ACPR) sequence (Nucleotides)**

ATGTATCTTTCGGCAGGATTGCGAAAAATTATGGATAATCCCCCCGTGACGTCGAAGTACCAGCACCGGAGTCAGGACGAGATGCTGTCGGCGAATTTGAATCACCAGCAAACGACAAATGGCAGTCCCTCGACGACGATGATGCTGGGTGGAGATGCGGCCGCTGTGGCATTTGCAAGTTCGTCGTCTAACGGAGCAGGGGATGCGTACTTCCGAGGATTGGCCGCACTGACGGGCAATGCTAGTGGGATCAGCTCAATGGACATGTCCAACAGCACCGAGACCGGCATTGTGGCACCAGGTCACACCGAAACGACCGTGGCCGTCATCATCGTGTACTGCGTGCTGTTCATCATCGCTGCCGGCGGAAATCTGTCCGTTGTGATAACCCTGTTTCGATCACGGCGCCATCGGAGGTCCCGAGTCAGTCTCATGATTTGCCATCTGGCCGTGGCGGACTTGATGGTGGCCTTCATCATGATCCCACTG**GAGGTCGGATGGCGCATTACGGTCCAGTGGCATGCGGGGAACGTGGCCTGCAAGGTG**TTCCTGTTCATGCGGGCGTTTTGTCTGTATCTGAGCTCGAATGTGTTGGTGTGTGTGTCGTTGGATCGGTGCTTTGCTGTGATATATCCGTTGCGGGTTTCGGCTGCCCGGAAGCGGGGCAAAATAATGCTCGGTGGAGCGTGGTTCATCGCATTCGCCAACGCCATTCCACAGAGTATAATCTTC**CGGGTTCAGCACCACCCAAACGTGCCGGACTTTACGCAGTGTGTGACGTTCGGGTTTTTCACCACCCCCGCCATGGAGACGGCTTACAAC**CTGTTCTGCGTGGTGGCAATGTACTTCATGCCGCTCATGGTCATCAGTGCAGCCTACACGGTTATCCTGTGCGAAATCTCCAATCGGTCCCGGGAAAAAGAGACGAGCGACACGAGCCACACCGGAGGGATGCGACTTCGTTGTAACGACTTGACGCACATCGAAAGGGCCCGGCAGCGGACACTCCGGCTCACCATTACCATCGTCGTCGTGTTCGTTTGGTGCTGGACGCCGTACGTTGTGATGACACTC**TGGTACATGTTTGACCGCGAAAGCGCCCTCAAAGTGGACGGTGCCATCCAGGAT**GGGCTCTTTCTGATGGCGGTGTCCAACTCATGCATGAACCCGCTGGTCTACGGTTCGTACGCGATGAAGTGTCGGAGGCCCTGGCGGAGGCAAATGGCACCAAATGGAGTGCAAACCCCAAACGCAGCCCAGAGGAGGTCCACGGATGCCGTATCGGGGATGGTCGGACCGCACTCGGATCGACTCACCGGGCGGGACAATAAGGACGAACTGGTGTACGAAGGCGGTGGCACCGAGCGGAACAAGCTGAAACAGTTCGGCATGGCCAACGGGATCTTCGCCAGGATTGGCAGCACCGGACGCAATCACGTTACAATGGCGGGTGGCGGCGCTGCCGGAACATCCGTGGCTACGACAAGGTTCGGAAACTGCACCACAACCGTAGGAGGGCGCAACGTTACCGAGCCAATCCGAACAGTGGCAATGAGCAGTGGCAGTAGAGTCAAAAGCTTCCAGCCGAGTTACAGTGAATTCGTTATGATGGCAGACGAAAGATCGCTGATGCTGTCGCGGCCTTCGTTCTACTCCGATCCAACTCCGAATGAGTTCGACTCGACAGTCGATTGTGGTCTCACTCGAAGTTTGGACACCGGTGGCTGGCGATGA

**Native *A. aegypti* ACPR** *(***ACPR) deduced sequence (Translation)**

MYLSAGLRKIMDNPPVTSKYQHRSQDEMLSANLNHQQTTNGSPSTTMMLGGDAAAVAFASSSSNGAGDAYFRGLAALTGNASGISSMDMSNSTETGIVAPGHTETTVAVIIVYCVLFIIAAGGNLSVVITLFRSRRHRRSRVSLMICHLAVADLMVAFIMIPL**EVGWRITVQWHAGNVACKV**FLFMRAFCLYLSSNVLVCVSLDRCFAVIYPLRVSAARKRGKIMLGGAWFIAFANAIPQSIIF**RVQHHPNVPDFTQCVTFGFFTTPAMETAYN**LFCVVAMYFMPLMVISAAYTVILCEISNRSREKETSDTSHTGGMRLRCNDLTHIERARQRTLRLTITIVVVFVWCWTPYVVMTL**WYMFDRESALKVDGAIQD**GLFLMAVSNSCMNPLVYGSYAMKCRRPWRRQMAPNGVQTPNAAQRRSTDAVSGMVGPHSDRLTGRDNKDELVYEGGGTERNKLKQFGMANGIFARIGSTGRNHVTMAGGGAAGTSVATTRFGNCTTTVGGRNVTEPIRTVAMSSGSRVKSFQPSYSEFVMMADERSLMLSRPSFYSDPTPNEFDSTVDCGLTRSLDTGGWR

**Native *A. aegypti* AKH receptor (AKHR) sequence (Nucleotides)**

AGTGCCCTAATGTACGGAGATCAAAGTGGTGTGAAACTTGRAAAAATGTCGGATTCTAGCATGCAAATCGCGTACAAGGATGAACACGCAACAGGATACTGATACTGCTTGATAAGTTAAATTGAAGGAGTTCAAAAAAACATATTTCAAAGCTTCCAACAAACAATTAGGTAACAACAGCTGAAAACACCTAAGATTCAAATTCGATTTCCACGGAACCATCCACTTTGGCTTTTTAACTTTGAACTAAAATGTCAAATGCAATTTTGAAAACAGAACGCGGCGAAGTGTTGAATTATAGTCACAGCTATGGAGAGAATTACAATAACGATGTGAACACTATGCCATACGTACTTTCGAGCTCTACGAGTAAAACCGGAGTTTTTGATAATGAAACATGGTACGGTACTAATAGTAGTAATTGGAATGAACCCTTGCCCATTGATATGCAATTCAACGATGGACATAAGTTGCAGATTGTTGTTTACAGCGTACTGATGGTCATTTCTGCCATAGGAAACATAACAGTGTTGGCACTTCTTATCAAACGGCGGCTCAAATCTCATTCCCGAATAGATATGATGTTAACCCATTTAGCTATAGCGGATTTATTGGTAACATTTCTAATGATGCCACTAGAGATTGGG**TGGGCGGCAACTGTCCAATGGAGGGCGGGCGATATCATGTGTCGT**GTGATGGCATTCTTCCGCACATTTGGATTACATCTATCAAGCTTCGTACTGGTCTGCATATCCGTAGATAGGTATTATGCTGTATTGCAGCCACTAAATCTTTCCAAAAGCAGAGGCAAAATAATGATATTGATTGCATGGGCAATGGCCACTCTCTGCAGTGCACCACAGCCCTTCATCTTCCATGTC**GAAATACACCCGAACCACACCTGGTATGAGCAATGTGTGACGTATAATACTTTCTCGAATGACAACTACCATACGGTC**TACAATATACTAGTAATGATGTTTATGTATGCATTGCCTTTGTTGACTATCATTTGTTCGTATGCGTCAATCTACATGGAAATCTTCAGACATAGTCGGATGCCAAACTCAGAGGGCTTTCGACGATCAAGCATTGATGTCCTAGGTCGCGCAAAGCGAAGAACATTGAAGATGACCATCACCATAGTTATGGCGTTTGTCATATGTTGGACCCCATATTATGTCATGTCTGTATGGTAT**TGGCTGGATCAAAAATCAGCGGAAAACGTTGATCAACGTGTG**CAAAAGGGTTTGTTTCTTTTTGCCTGTACCAACAGCTGCATGAACCCAATAGTGTACGGCATCTATAATGTGAAGCTTCGGAAAAAGAAAAAACCTGACGGTGTTAAGTCAGGCCAATCTTCCGTTATTCTAAGAAACTCTGCAAAATATACCAGGCACTCAGAATCCATCCGATCTTCATCAGGTAAAATCCGTGGCTACAGAGATCTTGATATACCGTCATGTGCAATCAAACCAGTCTGAAACCAAAGTTGAAATGTTGAAGCTGCTTACAAAATCGTAGTAG

**Native *A. aegypti* AKH receptor (AKHR) deduced sequence** (**Translation)**

MSNAILKTERGEVLNYSHSYGENYNNDVNTMPYVLSSSTSKTGVFDNETWYGTNSSNWNEPLPIDMQFNDGHKLQIVVYSVLMVISAIGNITVLALLIKRRLKSHSRIDMMLTHLAIADLLVTFLMMPLEIG**WAATVQWRAGDIMCR**VMAFFRTFGLHLSSFVLVCISVDRYYAVLQPLNLSKSRGKIMILIAWAMATLCSAPQPFIFHV**EIHPNHTWYEQCVTYNTFSNDNYHTV**YNILVMMFMYALPLLTIICSYASIYMEIFRHSRMPNSEGFRRSSIDVLGRAKRRTLKMTITIVMAFVICWTPYYVMSVWY**WLDQKSAENVDQRV**QKGLFLFACTNSCMNPIVYGIYNVKLRKKKKPDGVKSGQSSVILRNSAKYTRHSESIRSSSGKIRGYRDLDIPSCAIKPV

**ACPR ECL1 chimera (ACPR fullΔECL1) (Nucleotides)**

ATGTATCTTTCGGCAGGATTGCGAAAAATTATGGATAATCCCCCCGTGACGTCGAAGTACCAGCACCGGAGTCAGGACGAGATGCTGTCGGCGAATTTGAATCACCAGCAAACGACAAATGGCAGTCCCTCGACGACGATGATGCTGGGTGGAGATGCGGCCGCTGTGGCATTTGCAAGTTCGTCGTCTAACGGAGCAGGGGATGCGTACTTCCGAGGATTGGCCGCACTGACGGGCAATGCTAGTGGGATCAGCTCAATGGACATGTCCAACAGCACCGAGACCGGCATTGTGGCACCAGGTCACACCGAAACGACCGTGGCCGTCATCATCGTGTACTGCGTGCTGTTCATCATCGCTGCCGGCGGAAATCTGTCCGTTGTGATAACCCTGTTTCGATCACGGCGCCATCGGAGGTCCCGAGTCAGTCTCATGATTTGCCATCTGGCCGTGGCGGACTTGATGGTGGCCTTCATCATGATCCCACTG**TGGGCGGCAACTGTCCAATGGAGGGCGGGCGATATCATGTGTCGT**TTCCTGTTCATGCGGGCGTTTTGTCTGTATCTGAGCTCGAATGTGTTGGTGTGTGTGTCGTTGGATCGGTGCTTTGCTGTGATATATCCGTTGCGGGTTTCGGCTGCCCGGAAGCGGGGCAAAATAATGCTCGGTGGAGCGTGGTTCATCGCATTCGCCAACGCCATTCCACAGAGTATAATCTTCCGGGTTCAGCACCACCCAAACGTGCCGGACTTTACGCAGTGTGTGACGTTCGGGTTTTTCACCACCCCCGCCATGGAGACGGCTTACAACCTGTTCTGCGTGGTGGCAATGTACTTCATGCCGCTCATGGTCATCAGTGCAGCCTACACGGTTATCCTGTGCGAAATCTCCAATCGGTCCCGGGAAAAAGAGACGAGCGACACGAGCCACACCGGAGGGATGCGACTTCGTTGTAACGACTTGACGCACATCGAAAGGGCCCGGCAGCGGACACTCCGGCTCACCATTACCATCGTCGTCGTGTTCGTTTGGTGCTGGACGCCGTACGTTGTGATGACACTCTGGTACATGTTTGACCGCGAAAGCGCCCTCAAAGTGGACGGTGCCATCCAGGATGGGCTCTTTCTGATGGCGGTGTCCAACTCATGCATGAACCCGCTGGTCTACGGTTCGTACGCGATGAAGTGTCGGAGGCCCTGGCGGAGGCAAATGGCACCAAATGGAGTGCAAACCCCAAACGCAGCCCAGAGGAGGTCCACGGATGCCGTATCGGGGATGGTCGGACCGCACTCGGATCGACTCACCGGGCGGGACAATAAGGACGAACTGGTGTACGAAGGCGGTGGCACCGAGCGGAACAAGCTGAAACAGTTCGGCATGGCCAACGGGATCTTCGCCAGGATTGGCAGCACCGGACGCAATCACGTTACAATGGCGGGTGGCGGCGCTGCCGGAACATCCGTGGCTACGACAAGGTTCGGAAACTGCACCACAACCGTAGGAGGGCGCAACGTTACCGAGCCAATCCGAACAGTGGCAATGAGCAGTGGCAGTAGAGTCAAAAGCTTCCAGCCGAGTTACAGTGAATTCGTTATGATGGCAGACGAAAGATCGCTGATGCTGTCGCGGCCTTCGTTCTACTCCGATCCAACTCCGAATGAGTTCGACTCGACAGTCGATTGTGGTCTCACTCGAAGTTTGGACACCGGTGGCTGGCGATGA

**ACPR ECL1 chimera (ACPR fullΔECL1) deduced sequence (Translation)**

MYLSAGLRKIMDNPPVTSKYQHRSQDEMLSANLNHQQTTNGSPSTTMMLGGDAAAVAFASSSSNGAGDAYFRGLAALTGNASGISSMDMSNSTETGIVAPGHTETTVAVIIVYCVLFIIAAGGNLSVVITLFRSRRHRRSRVSLMICHLAVADLMVAFIMIPL**WAATVQWRAGDIMCR**FLFMRAFCLYLSSNVLVCVSLDRCFAVIYPLRVSAARKRGKIMLGGAWFIAFANAIPQSIIFRVQHHPNVPDFTQCVTFGFFTTPAMETAYNLFCVVAMYFMPLMVISAAYTVILCEISNRSREKETSDTSHTGGMRLRCNDLTHIERARQRTLRLTITIVVVFVWCWTPYVVMTLWYMFDRESALKVDGAIQDGLFLMAVSNSCMNPLVYGSYAMKCRRPWRRQMAPNGVQTPNAAQRRSTDAVSGMVGPHSDRLTGRDNKDELVYEGGGTERNKLKQFGMANGIFARIGSTGRNHVTMAGGGAAGTSVATTRFGNCTTTVGGRNVTEPIRTVAMSSGSRVKSFQPSYSEFVMMADERSLMLSRPSFYSDPTPNEFDSTVDCGLTRSLDTGGWR

**ACPR ECL2 chimera** (**ACPR fullΔECL 2) (Nucleotides)**

ATGTATCTTTCGGCAGGATTGCGAAAAATTATGGATAATCCCCCCGTGACGTCGAAGTACCAGCACCGGAGTCAGGACGAGATGCTGTCGGCGAATTTGAATCACCAGCAAACGACAAATGGCAGTCCCTCGACGACGATGATGCTGGGTGGAGATGCGGCCGCTGTGGCATTTGCAAGTTCGTCGTCTAACGGAGCAGGGGATGCGTACTTCCGAGGATTGGCCGCACTGACGGGCAATGCTAGTGGGATCAGCTCAATGGACATGTCCAACAGCACCGAGACCGGCATTGTGGCACCAGGTCACACCGAAACGACCGTGGCCGTCATCATCGTGTACTGCGTGCTGTTCATCATCGCTGCCGGCGGAAATCTGTCCGTTGTGATAACCCTGTTTCGATCACGGCGCCATCGGAGGTCCCGAGTCAGTCTCATGATTTGCCATCTGGCCGTGGCGGACTTGATGGTGGCCTTCATCATGATCCCACTGGAGGTCGGATGGCGCATTACGGTCCAGTGGCATGCGGGGAACGTGGCCTGCAAGGTGTTCCTGTTCATGCGGGCGTTTTGTCTGTATCTGAGCTCGAATGTGTTGGTGTGTGTGTCGTTGGATCGGTGCTTTGCTGTGATATATCCGTTGCGGGTTTCGGCTGCCCGGAAGCGGGGCAAAATAATGCTCGGTGGAGCGTGGTTCATCGCATTCGCCAACGCCATTCCACAGAGTATAATCTTC**GAAATACACCCGAACCACACCTGGTATGAGCAATGTGTGACGTATAATACTTTCTCGAATGACAACTACCATACGGTC**CTGTTCTGCGTGGTGGCAATGTACTTCATGCCGCTCATGGTCATCAGTGCAGCCTACACGGTTATCCTGTGCGAAATCTCCAATCGGTCCCGGGAAAAAGAGACGAGCGACACGAGCCACACCGGAGGGATGCGACTTCGTTGTAACGACTTGACGCACATCGAAAGGGCCCGGCAGCGGACACTCCGGCTCACCATTACCATCGTCGTCGTGTTCGTTTGGTGCTGGACGCCGTACGTTGTGATGACACTCTGGTACATGTTTGACCGCGAAAGCGCCCTCAAAGTGGACGGTGCCATCCAGGATGGGCTCTTTCTGATGGCGGTGTCCAACTCATGCATGAACCCGCTGGTCTACGGTTCGTACGCGATGAAGTGTCGGAGGCCCTGGCGGAGGCAAATGGCACCAAATGGAGTGCAAACCCCAAACGCAGCCCAGAGGAGGTCCACGGATGCCGTATCGGGGATGGTCGGACCGCACTCGGATCGACTCACCGGGCGGGACAATAAGGACGAACTGGTGTACGAAGGCGGTGGCACCGAGCGGAACAAGCTGAAACAGTTCGGCATGGCCAACGGGATCTTCGCCAGGATTGGCAGCACCGGACGCAATCACGTTACAATGGCGGGTGGCGGCGCTGCCGGAACATCCGTGGCTACGACAAGGTTCGGAAACTGCACCACAACCGTAGGAGGGCGCAACGTTACCGAGCCAATCCGAACAGTGGCAATGAGCAGTGGCAGTAGAGTCAAAAGCTTCCAGCCGAGTTACAGTGAATTCGTTATGATGGCAGACGAAAGATCGCTGATGCTGTCGCGGCCTTCGTTCTACTCCGATCCAACTCCGAATGAGTTCGACTCGACAGTCGATTGTGGTCTCACTCGAAGTTTGGACACCGGTGGCTGGCGATGA

**ACPR ECL2 chimera** **(ACPR fullΔECL 2) deduced sequence (Translation)**

MYLSAGLRKIMDNPPVTSKYQHRSQDEMLSANLNHQQTTNGSPSTTMMLGGDAAAVAFASSSSNGAGDAYFRGLAALTGNASGISSMDMSNSTETGIVAPGHTETTVAVIIVYCVLFIIAAGGNLSVVITLFRSRRHRRSRVSLMICHLAVADLMVAFIMIPLEVGWRITVQWHAGNVACKVFLFMRAFCLYLSSNVLVCVSLDRCFAVIYPLRVSAARKRGKIMLGGAWFIAFANAIPQSIIF**EIHPNHTWYEQCVTYNTFSNDNYHTV**LFCVVAMYFMPLMVISAAYTVILCEISNRSREKETSDTSHTGGMRLRCNDLTHIERARQRTLRLTITIVVVFVWCWTPYVVMTLWYMFDRESALKVDGAIQDGLFLMAVSNSCMNPLVYGSYAMKCRRPWRRQMAPNGVQTPNAAQRRSTDAVSGMVGPHSDRLTGRDNKDELVYEGGGTERNKLKQFGMANGIFARIGSTGRNHVTMAGGGAAGTSVATTRFGNCTTTVGGRNVTEPIRTVAMSSGSRVKSFQPSYSEFVMMADERSLMLSRPSFYSDPTPNEFDSTVDCGLTRSLDTGGWR

**ACPR ECL3 chimera (ACPR fullΔECL 3) (Nucleotides)**

ATGTATCTTTCGGCAGGATTGCGAAAAATTATGGATAATCCCCCCGTGACGTCGAAGTACCAGCACCGGAGTCAGGACGAGATGCTGTCGGCGAATTTGAATCACCAGCAAACGACAAATGGCAGTCCCTCGACGACGATGATGCTGGGTGGAGATGCGGCCGCTGTGGCATTTGCAAGTTCGTCGTCTAACGGAGCAGGGGATGCGTACTTCCGAGGATTGGCCGCACTGACGGGCAATGCTAGTGGGATCAGCTCAATGGACATGTCCAACAGCACCGAGACCGGCATTGTGGCACCAGGTCACACCGAAACGACCGTGGCCGTCATCATCGTGTACTGCGTGCTGTTCATCATCGCTGCCGGCGGAAATCTGTCCGTTGTGATAACCCTGTTTCGATCACGGCGCCATCGGAGGTCCCGAGTCAGTCTCATGATTTGCCATCTGGCCGTGGCGGACTTGATGGTGGCCTTCATCATGATCCCACTGGAGGTCGGATGGCGCATTACGGTCCAGTGGCATGCGGGGAACGTGGCCTGCAAGGTGTTCCTGTTCATGCGGGCGTTTTGTCTGTATCTGAGCTCGAATGTGTTGGTGTGTGTGTCGTTGGATCGGTGCTTTGCTGTGATATATCCGTTGCGGGTTTCGGCTGCCCGGAAGCGGGGCAAAATAATGCTCGGTGGAGCGTGGTTCATCGCATTCGCCAACGCCATTCCACAGAGTATAATCTTCCGGGTTCAGCACCACCCAAACGTGCCGGACTTTACGCAGTGTGTGACGTTCGGGTTTTTCACCACCCCCGCCATGGAGACGGCTTACAACCTGTTCTGCGTGGTGGCAATGTACTTCATGCCGCTCATGGTCATCAGTGCAGCCTACACGGTTATCCTGTGCGAAATCTCCAATCGGTCCCGGGAAAAAGAGACGAGCGACACGAGCCACACCGGAGGGATGCGACTTCGTTGTAACGACTTGACGCACATCGAAAGGGCCCGGCAGCGGACACTCCGGCTCACCATTACCATCGTCGTCGTGTTCGTTTGGTGCTGGACGCCGTACGTTGTGATGACACTC**TGGCTGGATCAAAAATCAGCGGAAAACGTTGATCAACGTGTG**GGGCTCTTTCTGATGGCGGTGTCCAACTCATGCATGAACCCGCTGGTCTACGGTTCGTACGCGATGAAGTGTCGGAGGCCCTGGCGGAGGCAAATGGCACCAAATGGAGTGCAAACCCCAAACGCAGCCCAGAGGAGGTCCACGGATGCCGTATCGGGGATGGTCGGACCGCACTCGGATCGACTCACCGGGCGGGACAATAAGGACGAACTGGTGTACGAAGGCGGTGGCACCGAGCGGAACAAGCTGAAACAGTTCGGCATGGCCAACGGGATCTTCGCCAGGATTGGCAGCACCGGACGCAATCACGTTACAATGGCGGGTGGCGGCGCTGCCGGAACATCCGTGGCTACGACAAGGTTCGGAAACTGCACCACAACCGTAGGAGGGCGCAACGTTACCGAGCCAATCCGAACAGTGGCAATGAGCAGTGGCAGTAGAGTCAAAAGCTTCCAGCCGAGTTACAGTGAATTCGTTATGATGGCAGACGAAAGATCGCTGATGCTGTCGCGGCCTTCGTTCTACTCCGATCCAACTCCGAATGAGTTCGACTCGACAGTCGATTGTGGTCTCACTCGAAGTTTGGACACCGGTGGCTGGCGATGA

**ACPR ECL3 chimera (ACPR fullΔECL 3) deduced sequence (Translation)**

MYLSAGLRKIMDNPPVTSKYQHRSQDEMLSANLNHQQTTNGSPSTTMMLGGDAAAVAFASSSSNGAGDAYFRGLAALTGNASGISSMDMSNSTETGIVAPGHTETTVAVIIVYCVLFIIAAGGNLSVVITLFRSRRHRRSRVSLMICHLAVADLMVAFIMIPLEVGWRITVQWHAGNVACKVFLFMRAFCLYLSSNVLVCVSLDRCFAVIYPLRVSAARKRGKIMLGGAWFIAFANAIPQSIIFRVQHHPNVPDFTQCVTFGFFTTPAMETAYNLFCVVAMYFMPLMVISAAYTVILCEISNRSREKETSDTSHTGGMRLRCNDLTHIERARQRTLRLTITIVVVFVWCWTPYVVMTL**WLDQKSAENVDQRV**GLFLMAVSNSCMNPLVYGSYAMKCRRPWRRQMAPNGVQTPNAAQRRSTDAVSGMVGPHSDRLTGRDNKDELVYEGGGTERNKLKQFGMANGIFARIGSTGRNHVTMAGGGAAGTSVATTRFGNCTTTVGGRNVTEPIRTVAMSSGSRVKSFQPSYSEFVMMADERSLMLSRPSFYSDPTPNEFDSTVDCGLTRSLDTGGWR

**Selectively modified residues within the ECL1 (ACPRΔECL1) (Nucleotides)**

ATGTATCTTTCGGCAGGATTGCGAAAAATTATGGATAATCCCCCCGTGACGTCGAAGTACCAGCACCGGAGTCAGGACGAGATGCTGTCGGCGAATTTGAATCACCAGCAAACGACAAATGGCAGTCCCTCGACGACGATGATGCTGGGTGGAGATGCGGCCGCTGTGGCATTTGCAAGTTCGTCGTCTAACGGAGCAGGGGATGCGTACTTCCGAGGATTGGCCGCACTGACGGGCAATGCTAGTGGGATCAGCTCAATGGACATGTCCAACAGCACCGAGACCGGCATTGTGGCACCAGGTCACACCGAAACGACCGTGGCCGTCATCATCGTGTACTGCGTGCTGTTCATCATCGCTGCCGGCGGAAATCTGTCCGTTGTGATAACCCTGTTTCGATCACGGCGCCATCGGAGGTCCCGAGTCAGTCTCATGATTTGCCATCTGGCCGTGGCGGACTTGATGGTGGCCTTCATCATGAT**GCCACTGGAGATCGGATGGGCCATTACGGTCCAGTGGCATGCGGGGGACGTGATGTGCCGGGTGATGCTGTTCTTC**CGGGCGTTTTGTCTGTATCTGAGCTCGAATGTGTTGGTGTGTGTGTCGTTGGATCGGTGCTTTGCTGTGATATATCCGTTGCGGGTTTCGGCTGCCCGGAAGCGGGGCAAAATAATGCTCGGTGGAGCGTGGTTCATCGCATTCGCCAACGCCATTCCACAGAGTATAATCTTCCGGGTTCAGCACCACCCAAACGTGCCGGACTTTACGCAGTGTGTGACGTTCGGGTTTTTCACCACCCCCGCCATGGAGACGGCTTACAACCTGTTCTGCGTGGTGGCAATGTACTTCATGCCGCTCATGGTCATCAGTGCAGCCTACACGGTTATCCTGTGCGAAATCTCCAATCGGTCCCGGGAAAAAGAGACGAGCGACACGAGCCACACCGGAGGGATGCGACTTCGTTGTAACGACTTGACGCACATCGAAAGGGCCCGGCAGCGGACACTCCGGCTCACCATTACCATCGTCGTCGTGTTCGTTTGGTGCTGGACGCCGTACGTTGTGATGACACTCTGGTACATGTTTGACCGCGAAAGCGCCCTCAAAGTGGACGGTGCCATCCAGGATGGGCTCTTTCTGATGGCGGTGTCCAACTCATGCATGAACCCGCTGGTCTACGGTTCGTACGCGATGAAGTGTCGGAGGCCCTGGCGGAGGCAAATGGCACCAAATGGAGTGCAAACCCCAAACGCAGCCCAGAGGAGGTCCACGGATGCCGTATCGGGGATGGTCGGACCGCACTCGGATCGACTCACCGGGCGGGACAATAAGGACGAACTGGTGTACGAAGGCGGTGGCACCGAGCGGAACAAGCTGAAACAGTTCGGCATGGCCAACGGGATCTTCGCCAGGATTGGCAGCACCGGACGCAATCACGTTACAATGGCGGGTGGCGGCGCTGCCGGAACATCCGTGGCTACGACAAGGTTCGGAAACTGCACCACAACCGTAGGAGGGCGCAACGTTACCGAGCCAATCCGAACAGTGGCAATGAGCAGTGGCAGTAGAGTCAAAAGCTTCCAGCCGAGTTACAGTGAATTCGTTATGATGGCAGACGAAAGATCGCTGATGCTGTCGCGGCCTTCGTTCTACTCCGATCCAACTCCGAATGAGTTCGACTCGACAGTCGATTGTGGTCTCACTCGAAGTTTGGACACCGGTGGCTGGCGATGA

**Selectively modified residues within the ECL1 (ACPRΔECL1) deduced sequence** **(Translation)**

MYLSAGLRKIMDNPPVTSKYQHRSQDEMLSANLNHQQTTNGSPSTTMMLGGDAAAVAFASSSSNGAGDAYFRGLAALTGNASGISSMDMSNSTETGIVAPGHTETTVAVIIVYCVLFIIAAGGNLSVVITLFRSRRHRRSRVSLMICHLAVADLMVAFIM**MPLEIGWAITVQWHAGDVMCRVMLFF**RAFCLYLSSNVLVCVSLDRCFAVIYPLRVSAARKRGKIMLGGAWFIAFANAIPQSIIFRVQHHPNVPDFTQCVTFGFFTTPAMETAYNLFCVVAMYFMPLMVISAAYTVILCEISNRSREKETSDTSHTGGMRLRCNDLTHIERARQRTLRLTITIVVVFVWCWTPYVVMTLWYMFDRESALKVDGAIQDGLFLMAVSNSCMNPLVYGSYAMKCRRPWRRQMAPNGVQTPNAAQRRSTDAVSGMVGPHSDRLTGRDNKDELVYEGGGTERNKLKQFGMANGIFARIGSTGRNHVTMAGGGAAGTSVATTRFGNCTTTVGGRNVTEPIRTVAMSSGSRVKSFQPSYSEFVMMADERSLMLSRPSFYSDPTPNEFDSTVDCGLTRSLDTGGWR

**Selectively modified residues within the ECL2 (ACPRΔECL2) (Nucleotides)**

ATGTATCTTTCGGCAGGATTGCGAAAAATTATGGATAATCCCCCCGTGACGTCGAAGTACCAGCACCGGAGTCAGGACGAGATGCTGTCGGCGAATTTGAATCACCAGCAAACGACAAATGGCAGTCCCTCGACGACGATGATGCTGGGTGGAGATGCGGCCGCTGTGGCATTTGCAAGTTCGTCGTCTAACGGAGCAGGGGATGCGTACTTCCGAGGATTGGCCGCACTGACGGGCAATGCTAGTGGGATCAGCTCAATGGACATGTCCAACAGCACCGAGACCGGCATTGTGGCACCAGGTCACACCGAAACGACCGTGGCCGTCATCATCGTGTACTGCGTGCTGTTCATCATCGCTGCCGGCGGAAATCTGTCCGTTGTGATAACCCTGTTTCGATCACGGCGCCATCGGAGGTCCCGAGTCAGTCTCATGATTTGCCATCTGGCCGTGGCGGACTTGATGGTGGCCTTCATCATGATCCCACTGGAGGTCGGATGGCGCATTACGGTCCAGTGGCATGCGGGGAACGTGGCCTGCAAGGTGTTCCTGTTCATGCGGGCGTTTTGTCTGTATCTGAGCTCGAATGTGTTGGTGTGTGTGTCGTTGGATCGGTGCTTTGCTGTGATATATCCGTTGCGGGTTTCGGCTGCCCGGAAGCGGGGCAAAATAATGCTCGGTGGAGCGTGGTTCATCGCATTCGCCAACGCCATTCCACAGAGTATAATCTTC**CACGTTGAGCACCACCCAAACGTGACGGACTATGAGCAGTGTGTGACGTTCAACTTTTTCACCACCCCCGCCATGGAGACGGCTTACAACCTGTTCGGCA**TGGTGGCAATGTACTTCATGCCGCTCATGGTCATCAGTGCAGCCTACACGGTTATCCTGTGCGAAATCTCCAATCGGTCCCGGGAAAAAGAGACGAGCGACACGAGCCACACCGGAGGGATGCGACTTCGTTGTAACGACTTGACGCACATCGAAAGGGCCCGGCAGCGGACACTCCGGCTCACCATTACCATCGTCGTCGTGTTCGTTTGGTGCTGGACGCCGTACGTTGTGATGACACTCTGGTACATGTTTGACCGCGAAAGCGCCCTCAAAGTGGACGGTGCCATCCAGGATGGGCTCTTTCTGATGGCGGTGTCCAACTCATGCATGAACCCGCTGGTCTACGGTTCGTACGCGATGAAGTGTCGGAGGCCCTGGCGGAGGCAAATGGCACCAAATGGAGTGCAAACCCCAAACGCAGCCCAGAGGAGGTCCACGGATGCCGTATCGGGGATGGTCGGACCGCACTCGGATCGACTCACCGGGCGGGACAATAAGGACGAACTGGTGTACGAAGGCGGTGGCACCGAGCGGAACAAGCTGAAACAGTTCGGCATGGCCAACGGGATCTTCGCCAGGATTGGCAGCACCGGACGCAATCACGTTACAATGGCGGGTGGCGGCGCTGCCGGAACATCCGTGGCTACGACAAGGTTCGGAAACTGCACCACAACCGTAGGAGGGCGCAACGTTACCGAGCCAATCCGAACAGTGGCAATGAGCAGTGGCAGTAGAGTCAAAAGCTTCCAGCCGAGTTACAGTGAATTCGTTATGATGGCAGACGAAAGATCGCTGATGCTGTCGCGGCCTTCGTTCTACTCCGATCCAACTCCGAATGAGTTCGACTCGACAGTCGATTGTGGTCTCACTCGAAGTTTGGACACCGGTGGCTGGCGATGA

**Selectively modified residues within the ECL2 (ACPRΔECL2) deduced sequence** **(Translation)**

MYLSAGLRKIMDNPPVTSKYQHRSQDEMLSANLNHQQTTNGSPSTTMMLGGDAAAVAFASSSSNGAGDAYFRGLAALTGNASGISSMDMSNSTETGIVAPGHTETTVAVIIVYCVLFIIAAGGNLSVVITLFRSRRHRRSRVSLMICHLAVADLMVAFIMIPLEVGWRITVQWHAGNVACKVFLFMRAFCLYLSSNVLVCVSLDRCFAVIYPLRVSAARKRGKIMLGGAWFIAFANAIPQSIIF**HVEHHPNVTDYEQCVTFNFFTTPAMETAYNLFGM**VAMYFMPLMVISAAYTVILCEISNRSREKETSDTSHTGGMRLRCNDLTHIERARQRTLRLTITIVVVFVWCWTPYVVMTLWYMFDRESALKVDGAIQDGLFLMAVSNSCMNPLVYGSYAMKCRRPWRRQMAPNGVQTPNAAQRRSTDAVSGMVGPHSDRLTGRDNKDELVYEGGGTERNKLKQFGMANGIFARIGSTGRNHVTMAGGGAAGTSVATTRFGNCTTTVGGRNVTEPIRTVAMSSGSRVKSFQPSYSEFVMMADERSLMLSRPSFYSDPTPNEFDSTVDCGLTRSLDTGGWR

**Selectively modified residues within the ECL3 (ACPRΔECL3) (Nucleotides)**

ATGTATCTTTCGGCAGGATTGCGAAAAATTATGGATAATCCCCCCGTGACGTCGAAGTACCAGCACCGGAGTCAGGACGAGATGCTGTCGGCGAATTTGAATCACCAGCAAACGACAAATGGCAGTCCCTCGACGACGATGATGCTGGGTGGAGATGCGGCCGCTGTGGCATTTGCAAGTTCGTCGTCTAACGGAGCAGGGGATGCGTACTTCCGAGGATTGGCCGCACTGACGGGCAATGCTAGTGGGATCAGCTCAATGGACATGTCCAACAGCACCGAGACCGGCATTGTGGCACCAGGTCACACCGAAACGACCGTGGCCGTCATCATCGTGTACTGCGTGCTGTTCATCATCGCTGCCGGCGGAAATCTGTCCGTTGTGATAACCCTGTTTCGATCACGGCGCCATCGGAGGTCCCGAGTCAGTCTCATGATTTGCCATCTGGCCGTGGCGGACTTGATGGTGGCCTTCATCATGATCCCACTGGAGGTCGGATGGCGCATTACGGTCCAGTGGCATGCGGGGAACGTGGCCTGCAAGGTGTTCCTGTTCATGCGGGCGTTTTGTCTGTATCTGAGCTCGAATGTGTTGGTGTGTGTGTCGTTGGATCGGTGCTTTGCTGTGATATATCCGTTGCGGGTTTCGGCTGCCCGGAAGCGGGGCAAAATAATGCTCGGTGGAGCGTGGTTCATCGCATTCGCCAACGCCATTCCACAGAGTATAATCTTCCGGGTTCAGCACCACCCAAACGTGCCGGACTTTACGCAGTGTGTGACGTTCGGGTTTTTCACCACCCCCGCCATGGAGACGGCTTACAACCTGTTCTGCGTGGTGGCAATGTACTTCATGCCGCTCATGGTCATCAGTGCAGCCTACACGGTTATCCTGTGCGAAATCTCCAATCGGTCCCGGGAAAAAGAGACGAGCGACACGAGCCACACCGGAGGGATGCGACTTCGTTGTAACGACTTGACGCACATCGAAAGGGCCCGGCAGCGGACACTCCGGCTCACCATTACCATCGTCGTCGTGTTCGTTTGGTGCTGGACGCCGTAC**TATGTGATGTCACTCTGGTACTGGCTTGACCGCAAAAGCGCCCTCAAAGTGGACCAGCGCATCCAGAAAGGGCTCTTTCTGTTC**GCGGTGTCCAACTCATGCATGAACCCGCTGGTCTACGGTTCGTACGCGATGAAGTGTCGGAGGCCCTGGCGGAGGCAAATGGCACCAAATGGAGTGCAAACCCCAAACGCAGCCCAGAGGAGGTCCACGGATGCCGTATCGGGGATGGTCGGACCGCACTCGGATCGACTCACCGGGCGGGACAATAAGGACGAACTGGTGTACGAAGGCGGTGGCACCGAGCGGAACAAGCTGAAACAGTTCGGCATGGCCAACGGGATCTTCGCCAGGATTGGCAGCACCGGACGCAATCACGTTACAATGGCGGGTGGCGGCGCTGCCGGAACATCCGTGGCTACGACAAGGTTCGGAAACTGCACCACAACCGTAGGAGGGCGCAACGTTACCGAGCCAATCCGAACAGTGGCAATGAGCAGTGGCAGTAGAGTCAAAAGCTTCCAGCCGAGTTACAGTGAATTCGTTATGATGGCAGACGAAAGATCGCTGATGCTGTCGCGGCCTTCGTTCTACTCCGATCCAACTCCGAATGAGTTCGACTCGACAGTCGATTGTGGTCTCACTCGAAGTTTGGACACCGGTGGCTGGCGATGA

**Selectively modified residues within the ECL2 (ACPRΔECL3) deduced sequence** **(Translation)**

MYLSAGLRKIMDNPPVTSKYQHRSQDEMLSANLNHQQTTNGSPSTTMMLGGDAAAVAFASSSSNGAGDAYFRGLAALTGNASGISSMDMSNSTETGIVAPGHTETTVAVIIVYCVLFIIAAGGNLSVVITLFRSRRHRRSRVSLMICHLAVADLMVAFIMIPLEVGWRITVQWHAGNVACKVFLFMRAFCLYLSSNVLVCVSLDRCFAVIYPLRVSAARKRGKIMLGGAWFIAFANAIPQSIIFRVQHHPNVPDFTQCVTFGFFTTPAMETAYNLFCVVAMYFMPLMVISAAYTVILCEISNRSREKETSDTSHTGGMRLRCNDLTHIERARQRTLRLTITIVVVFVWCWTPY**YVMSLWYWLDRKSALKVDQRIQKGLFLF**AVSNSCMNPLVYGSYAMKCRRPWRRQMAPNGVQTPNAAQRRSTDAVSGMVGPHSDRLTGRDNKDELVYEGGGTERNKLKQFGMANGIFARIGSTGRNHVTMAGGGAAGTSVATTRFGNCTTTVGGRNVTEPIRTVAMSSGSRVKSFQPSYSEFVMMADERSLMLSRPSFYSDPTPNEFDSTVDCGLTRSLDTGGWR

**ACPR ECL1 chimera with ACPR ECL2 chimera** (**ACPRΔECL 1+2)**

ATGTATCTTTCGGCAGGATTGCGAAAAATTATGGATAATCCCCCCGTGACGTCGAAGTACCAGCACCGGAGTCAGGACGAGATGCTGTCGGCGAATTTGAATCACCAGCAAACGACAAATGGCAGTCCCTCGACGACGATGATGCTGGGTGGAGATGCGGCCGCTGTGGCATTTGCAAGTTCGTCGTCTAACGGAGCAGGGGATGCGTACTTCCGAGGATTGGCCGCACTGACGGGCAATGCTAGTGGGATCAGCTCAATGGACATGTCCAACAGCACCGAGACCGGCATTGTGGCACCAGGTCACACCGAAACGACCGTGGCCGTCATCATCGTGTACTGCGTGCTGTTCATCATCGCTGCCGGCGGAAATCTGTCCGTTGTGATAACCCTGTTTCGATCACGGCGCCATCGGAGGTCCCGAGTCAGTCTCATGATTTGCCATCTGGCCGTGGCGGACTTGATGGTGGCCTTCATCATGAT**GCCACTGGAGATCGGATGGGCCATTACGGTCCAGTGGCATGCGGGGGACGTGATGTGCCGGGTGATGCTGTTCTTC**CGGGCGTTTTGTCTGTATCTGAGCTCGAATGTGTTGGTGTGTGTGTCGTTGGATCGGTGCTTTGCTGTGATATATCCGTTGCGGGTTTCGGCTGCCCGGAAGCGGGGCAAAATAATGCTCGGTGGAGCGTGGTTCATCGCATTCGCCAACGCCATTCCACAGAGTATAATCTTC**CACGTTGAGCACCACCCAAACGTGACGGACTATGAGCAGTGTGTGACGTTCAACTTTTTCACCACCCCCGCCATGGAGACGGCTTACAACCTGTTCGGCA**TGGTGGCAATGTACTTCATGCCGCTCATGGTCATCAGTGCAGCCTACACGGTTATCCTGTGCGAAATCTCCAATCGGTCCCGGGAAAAAGAGACGAGCGACACGAGCCACACCGGAGGGATGCGACTTCGTTGTAACGACTTGACGCACATCGAAAGGGCCCGGCAGCGGACACTCCGGCTCACCATTACCATCGTCGTCGTGTTCGTTTGGTGCTGGACGCCGTACGTTGTGATGACACTCTGGTACATGTTTGACCGCGAAAGCGCCCTCAAAGTGGACGGTGCCATCCAGGATGGGCTCTTTCTGATGGCGGTGTCCAACTCATGCATGAACCCGCTGGTCTACGGTTCGTACGCGATGAAGTGTCGGAGGCCCTGGCGGAGGCAAATGGCACCAAATGGAGTGCAAACCCCAAACGCAGCCCAGAGGAGGTCCACGGATGCCGTATCGGGGATGGTCGGACCGCACTCGGATCGACTCACCGGGCGGGACAATAAGGACGAACTGGTGTACGAAGGCGGTGGCACCGAGCGGAACAAGCTGAAACAGTTCGGCATGGCCAACGGGATCTTCGCCAGGATTGGCAGCACCGGACGCAATCACGTTACAATGGCGGGTGGCGGCGCTGCCGGAACATCCGTGGCTACGACAAGGTTCGGAAACTGCACCACAACCGTAGGAGGGCGCAACGTTACCGAGCCAATCCGAACAGTGGCAATGAGCAGTGGCAGTAGAGTCAAAAGCTTCCAGCCGAGTTACAGTGAATTCGTTATGATGGCAGACGAAAGATCGCTGATGCTGTCGCGGCCTTCGTTCTACTCCGATCCAACTCCGAATGAGTTCGACTCGACAGTCGATTGTGGTCTCACTCGAAGTTTGGACACCGGTGGCTGGCGATGA

**ACPR ECL1 chimera with ACPR ECL2 chimera** (**ACPRΔECL 1+2) deduced sequence** **(Translation)**

MYLSAGLRKIMDNPPVTSKYQHRSQDEMLSANLNHQQTTNGSPSTTMMLGGDAAAVAFASSSSNGAGDAYFRGLAALTGNASGISSMDMSNSTETGIVAPGHTETTVAVIIVYCVLFIIAAGGNLSVVITLFRSRRHRRSRVSLMICHLAVADLMVAFIM**MPLEIGWAITVQWHAGDVMCRVMLFF**RAFCLYLSSNVLVCVSLDRCFAVIYPLRVSAARKRGKIMLGGAWFIAFANAIPQSIIF**HVEHHPNVTDYEQCVTFNFFTTPAMETAYNLFGM**VAMYFMPLMVISAAYTVILCEISNRSREKETSDTSHTGGMRLRCNDLTHIERARQRTLRLTITIVVVFVWCWTPYVVMTLWYMFDRESALKVDGAIQDGLFLMAVSNSCMNPLVYGSYAMKCRRPWRRQMAPNGVQTPNAAQRRSTDAVSGMVGPHSDRLTGRDNKDELVYEGGGTERNKLKQFGMANGIFARIGSTGRNHVTMAGGGAAGTSVATTRFGNCTTTVGGRNVTEPIRTVAMSSGSRVKSFQPSYSEFVMMADERSLMLSRPSFYSDPTPNEFDSTVDCGLTRSLDTGGWR

**ACPR ECL2 chimera with ACPR ECL3 chimera** (**ACPRΔECL 2+3) (Nucleotides)**

ATGTATCTTTCGGCAGGATTGCGAAAAATTATGGATAATCCCCCCGTGACGTCGAAGTACCAGCACCGGAGTCAGGACGAGATGCTGTCGGCGAATTTGAATCACCAGCAAACGACAAATGGCAGTCCCTCGACGACGATGATGCTGGGTGGAGATGCGGCCGCTGTGGCATTTGCAAGTTCGTCGTCTAACGGAGCAGGGGATGCGTACTTCCGAGGATTGGCCGCACTGACGGGCAATGCTAGTGGGATCAGCTCAATGGACATGTCCAACAGCACCGAGACCGGCATTGTGGCACCAGGTCACACCGAAACGACCGTGGCCGTCATCATCGTGTACTGCGTGCTGTTCATCATCGCTGCCGGCGGAAATCTGTCCGTTGTGATAACCCTGTTTCGATCACGGCGCCATCGGAGGTCCCGAGTCAGTCTCATGATTTGCCATCTGGCCGTGGCGGACTTGATGGTGGCCTTCATCATGATCCCACTGGAGGTCGGATGGCGCATTACGGTCCAGTGGCATGCGGGGAACGTGGCCTGCAAGGTGTTCCTGTTCATGCGGGCGTTTTGTCTGTATCTGAGCTCGAATGTGTTGGTGTGTGTGTCGTTGGATCGGTGCTTTGCTGTGATATATCCGTTGCGGGTTTCGGCTGCCCGGAAGCGGGGCAAAATAATGCTCGGTGGAGCGTGGTTCATCGCATTCGCCAACGCCATTCCACAGAGTATAATCTTC**CACGTTGAGCACCACCCAAACGTGACGGACTATGAGCAGTGTGTGACGTTCAACTTTTTCACCACCCCCGCCATGGAGACGGCTTACAACCTGTTCGGCA**TGGTGGCAATGTACTTCATGCCGCTCATGGTCATCAGTGCAGCCTACACGGTTATCCTGTGCGAAATCTCCAATCGGTCCCGGGAAAAAGAGACGAGCGACACGAGCCACACCGGAGGGATGCGACTTCGTTGTAACGACTTGACGCACATCGAAAGGGCCCGGCAGCGGACACTCCGGCTCACCATTACCATCGTCGTCGTGTTCGTTTGGTGCTGGACGCCGTAC**TATGTGATGTCACTCTGGTACTGGCTTGACCGCAAAAGCGCCCTCAAAGTGGACCAGCGCATCCAGAAAGGGCTCTTTCTGTTC**GCGGTGTCCAACTCATGCATGAACCCGCTGGTCTACGGTTCGTACGCGATGAAGTGTCGGAGGCCCTGGCGGAGGCAAATGGCACCAAATGGAGTGCAAACCCCAAACGCAGCCCAGAGGAGGTCCACGGATGCCGTATCGGGGATGGTCGGACCGCACTCGGATCGACTCACCGGGCGGGACAATAAGGACGAACTGGTGTACGAAGGCGGTGGCACCGAGCGGAACAAGCTGAAACAGTTCGGCATGGCCAACGGGATCTTCGCCAGGATTGGCAGCACCGGACGCAATCACGTTACAATGGCGGGTGGCGGCGCTGCCGGAACATCCGTGGCTACGACAAGGTTCGGAAACTGCACCACAACCGTAGGAGGGCGCAACGTTACCGAGCCAATCCGAACAGTGGCAATGAGCAGTGGCAGTAGAGTCAAAAGCTTCCAGCCGAGTTACAGTGAATTCGTTATGATGGCAGACGAAAGATCGCTGATGCTGTCGCGGCCTTCGTTCTACTCCGATCCAACTCCGAATGAGTTCGACTCGACAGTCGATTGTGGTCTCACTCGAAGTTTGGACACCGGTGGCTGGCGATGA

**ACPR ECL2 chimera with ACPR ECL3 chimera** (**ACPRΔECL 2+3) deduced sequence** **(Translation)**

MYLSAGLRKIMDNPPVTSKYQHRSQDEMLSANLNHQQTTNGSPSTTMMLGGDAAAVAFASSSSNGAGDAYFRGLAALTGNASGISSMDMSNSTETGIVAPGHTETTVAVIIVYCVLFIIAAGGNLSVVITLFRSRRHRRSRVSLMICHLAVADLMVAFIMIPLEVGWRITVQWHAGNVACKVFLFMRAFCLYLSSNVLVCVSLDRCFAVIYPLRVSAARKRGKIMLGGAWFIAFANAIPQSIIF**HVEHHPNVTDYEQCVTFNFFTTPAMETAYNLFGM**VAMYFMPLMVISAAYTVILCEISNRSREKETSDTSHTGGMRLRCNDLTHIERARQRTLRLTITIVVVFVWCWTPY**YVMSLWYWLDRKSALKVDQRIQKGLFLF**AVSNSCMNPLVYGSYAMKCRRPWRRQMAPNGVQTPNAAQRRSTDAVSGMVGPHSDRLTGRDNKDELVYEGGGTERNKLKQFGMANGIFARIGSTGRNHVTMAGGGAAGTSVATTRFGNCTTTVGGRNVTEPIRTVAMSSGSRVKSFQPSYSEFVMMADERSLMLSRPSFYSDPTPNEFDSTVDCGLTRSLDTGGWR

**ACPR ECL1 chimera with ACPR ECL3 chimera** (**ACPRΔECL 1+3) (Nucleotides)**

ATGTATCTTTCGGCAGGATTGCGAAAAATTATGGATAATCCCCCCGTGACGTCGAAGTACCAGCACCGGAGTCAGGACGAGATGCTGTCGGCGAATTTGAATCACCAGCAAACGACAAATGGCAGTCCCTCGACGACGATGATGCTGGGTGGAGATGCGGCCGCTGTGGCATTTGCAAGTTCGTCGTCTAACGGAGCAGGGGATGCGTACTTCCGAGGATTGGCCGCACTGACGGGCAATGCTAGTGGGATCAGCTCAATGGACATGTCCAACAGCACCGAGACCGGCATTGTGGCACCAGGTCACACCGAAACGACCGTGGCCGTCATCATCGTGTACTGCGTGCTGTTCATCATCGCTGCCGGCGGAAATCTGTCCGTTGTGATAACCCTGTTTCGATCACGGCGCCATCGGAGGTCCCGAGTCAGTCTCATGATTTGCCATCTGGCCGTGGCGGACTTGATGGTGGCCTTCATCATGAT**GCCACTGGAGATCGGATGGGCCATTACGGTCCAGTGGCATGCGGGGGACGTGATGTGCCGGGTGATGCTGTTCTTC**CGGGCGTTTTGTCTGTATCTGAGCTCGAATGTGTTGGTGTGTGTGTCGTTGGATCGGTGCTTTGCTGTGATATATCCGTTGCGGGTTTCGGCTGCCCGGAAGCGGGGCAAAATAATGCTCGGTGGAGCGTGGTTCATCGCATTCGCCAACGCCATTCCACAGAGTATAATCTTCCGGGTTCAGCACCACCCAAACGTGCCGGACTTTACGCAGTGTGTGACGTTCGGGTTTTTCACCACCCCCGCCATGGAGACGGCTTACAACCTGTTCTGCGTGGTGGCAATGTACTTCATGCCGCTCATGGTCATCAGTGCAGCCTACACGGTTATCCTGTGCGAAATCTCCAATCGGTCCCGGGAAAAAGAGACGAGCGACACGAGCCACACCGGAGGGATGCGACTTCGTTGTAACGACTTGACGCACATCGAAAGGGCCCGGCAGCGGACACTCCGGCTCACCATTACCATCGTCGTCGTGTTCGTTTGGTGCTGGACGCCGTAC**TATGTGATGTCACTCTGGTACTGGCTTGACCGCAAAAGCGCCCTCAAAGTGGACCAGCGCATCCAGAAAGGGCTCTTTCTGTTC**GCGGTGTCCAACTCATGCATGAACCCGCTGGTCTACGGTTCGTACGCGATGAAGTGTCGGAGGCCCTGGCGGAGGCAAATGGCACCAAATGGAGTGCAAACCCCAAACGCAGCCCAGAGGAGGTCCACGGATGCCGTATCGGGGATGGTCGGACCGCACTCGGATCGACTCACCGGGCGGGACAATAAGGACGAACTGGTGTACGAAGGCGGTGGCACCGAGCGGAACAAGCTGAAACAGTTCGGCATGGCCAACGGGATCTTCGCCAGGATTGGCAGCACCGGACGCAATCACGTTACAATGGCGGGTGGCGGCGCTGCCGGAACATCCGTGGCTACGACAAGGTTCGGAAACTGCACCACAACCGTAGGAGGGCGCAACGTTACCGAGCCAATCCGAACAGTGGCAATGAGCAGTGGCAGTAGAGTCAAAAGCTTCCAGCCGAGTTACAGTGAATTCGTTATGATGGCAGACGAAAGATCGCTGATGCTGTCGCGGCCTTCGTTCTACTCCGATCCAACTCCGAATGAGTTCGACTCGACAGTCGATTGTGGTCTCACTCGAAGTTTGGACACCGGTGGCTGGCGATGA

**ACPR ECL1 chimera with ACPR ECL3 chimera** (**ACPRΔECL 1+3) deduced sequence** **(Translation)**

MYLSAGLRKIMDNPPVTSKYQHRSQDEMLSANLNHQQTTNGSPSTTMMLGGDAAAVAFASSSSNGAGDAYFRGLAALTGNASGISSMDMSNSTETGIVAPGHTETTVAVIIVYCVLFIIAAGGNLSVVITLFRSRRHRRSRVSLMICHLAVADLMVAFIM**MPLEIGWAITVQWHAGDVMCRVMLFF**RAFCLYLSSNVLVCVSLDRCFAVIYPLRVSAARKRGKIMLGGAWFIAFANAIPQSIIFRVQHHPNVPDFTQCVTFGFFTTPAMETAYNLFCVVAMYFMPLMVISAAYTVILCEISNRSREKETSDTSHTGGMRLRCNDLTHIERARQRTLRLTITIVVVFVWCWTPY**YVMSLWYWLDRKSALKVDQRIQKGLFLF**AVSNSCMNPLVYGSYAMKCRRPWRRQMAPNGVQTPNAAQRRSTDAVSGMVGPHSDRLTGRDNKDELVYEGGGTERNKLKQFGMANGIFARIGSTGRNHVTMAGGGAAGTSVATTRFGNCTTTVGGRNVTEPIRTVAMSSGSRVKSFQPSYSEFVMMADERSLMLSRPSFYSDPTPNEFDSTVDCGLTRSLDTGGWR

**Three selectively modified ACPR ECLs chimeras** (**ACPRΔECL1+2+3) (Nucleotides)**

ATGTATCTTTCGGCAGGATTGCGAAAAATTATGGATAATCCCCCCGTGACGTCGAAGTACCAGCACCGGAGTCAGGACGAGATGCTGTCGGCGAATTTGAATCACCAGCAAACGACAAATGGCAGTCCCTCGACGACGATGATGCTGGGTGGAGATGCGGCCGCTGTGGCATTTGCAAGTTCGTCGTCTAACGGAGCAGGGGATGCGTACTTCCGAGGATTGGCCGCACTGACGGGCAATGCTAGTGGGATCAGCTCAATGGACATGTCCAACAGCACCGAGACCGGCATTGTGGCACCAGGTCACACCGAAACGACCGTGGCCGTCATCATCGTGTACTGCGTGCTGTTCATCATCGCTGCCGGCGGAAATCTGTCCGTTGTGATAACCCTGTTTCGATCACGGCGCCATCGGAGGTCCCGAGTCAGTCTCATGATTTGCCATCTGGCCGTGGCGGACTTGATGGTGGCCTTCATCATGAT**GCCACTGGAGATCGGATGGGCCATTACGGTCCAGTGGCATGCGGGGGACGTGATGTGCCGGGTGATGCTGTTCTTC**CGGGCGTTTTGTCTGTATCTGAGCTCGAATGTGTTGGTGTGTGTGTCGTTGGATCGGTGCTTTGCTGTGATATATCCGTTGCGGGTTTCGGCTGCCCGGAAGCGGGGCAAAATAATGCTCGGTGGAGCGTGGTTCATCGCATTCGCCAACGCCATTCCACAGAGTATAATCTTC**CACGTTGAGCACCACCCAAACGTGACGGACTATGAGCAGTGTGTGACGTTCAACTTTTTCACCACCCCCGCCATGGAGACGGCTTACAACCTGTTCGGCA**TGGTGGCAATGTACTTCATGCCGCTCATGGTCATCAGTGCAGCCTACACGGTTATCCTGTGCGAAATCTCCAATCGGTCCCGGGAAAAAGAGACGAGCGACACGAGCCACACCGGAGGGATGCGACTTCGTTGTAACGACTTGACGCACATCGAAAGGGCCCGGCAGCGGACACTCCGGCTCACCATTACCATCGTCGTCGTGTTCGTTTGGTGCTGGACGCCGTAC**TATGTGATGTCACTCTGGTACTGGCTTGACCGCAAAAGCGCCCTCAAAGTGGACCAGCGCATCCAGAAAGGGCTCTTTCTGTTC**GCGGTGTCCAACTCATGCATGAACCCGCTGGTCTACGGTTCGTACGCGATGAAGTGTCGGAGGCCCTGGCGGAGGCAAATGGCACCAAATGGAGTGCAAACCCCAAACGCAGCCCAGAGGAGGTCCACGGATGCCGTATCGGGGATGGTCGGACCGCACTCGGATCGACTCACCGGGCGGGACAATAAGGACGAACTGGTGTACGAAGGCGGTGGCACCGAGCGGAACAAGCTGAAACAGTTCGGCATGGCCAACGGGATCTTCGCCAGGATTGGCAGCACCGGACGCAATCACGTTACAATGGCGGGTGGCGGCGCTGCCGGAACATCCGTGGCTACGACAAGGTTCGGAAACTGCACCACAACCGTAGGAGGGCGCAACGTTACCGAGCCAATCCGAACAGTGGCAATGAGCAGTGGCAGTAGAGTCAAAAGCTTCCAGCCGAGTTACAGTGAATTCGTTATGATGGCAGACGAAAGATCGCTGATGCTGTCGCGGCCTTCGTTCTACTCCGATCCAACTCCGAATGAGTTCGACTCGACAGTCGATTGTGGTCTCACTCGAAGTTTGGACACCGGTGGCTGGCGATGA

**Three selectively modified ACPR ECLs chimeras (ACPRΔECL1+2+3) deduced sequence** **(Translation)**

MYLSAGLRKIMDNPPVTSKYQHRSQDEMLSANLNHQQTTNGSPSTTMMLGGDAAAVAFASSSSNGAGDAYFRGLAALTGNASGISSMDMSNSTETGIVAPGHTETTVAVIIVYCVLFIIAAGGNLSVVITLFRSRRHRRSRVSLMICHLAVADLMVAFIM**MPLEIGWAITVQWHAGDVMCRVMLFF**RAFCLYLSSNVLVCVSLDRCFAVIYPLRVSAARKRGKIMLGGAWFIAFANAIPQSIIF**HVEHHPNVTDYEQCVTFNFFTTPAMETAYNLFGM**VAMYFMPLMVISAAYTVILCEISNRSREKETSDTSHTGGMRLRCNDLTHIERARQRTLRLTITIVVVFVWCWTPY**YVMSLWYWLDRKSALKVDQRIQKGLFLF**AVSNSCMNPLVYGSYAMKCRRPWRRQMAPNGVQTPNAAQRRSTDAVSGMVGPHSDRLTGRDNKDELVYEGGGTERNKLKQFGMANGIFARIGSTGRNHVTMAGGGAAGTSVATTRFGNCTTTVGGRNVTEPIRTVAMSSGSRVKSFQPSYSEFVMMADERSLMLSRPSFYSDPTPNEFDSTVDCGLTRSLDTGGWR
